# Posterior hippocampal spindle-ripples co-occur with neocortical theta-bursts and down-upstates, and phase-lock with parietal spindles during NREM sleep in humans

**DOI:** 10.1101/702936

**Authors:** Xi Jiang, Jorge Gonzalez-Martinez, Eric Halgren

**Affiliations:** Department of Neurosciences, University of California at San Diego, La Jolla, CA 92093, USA; Department of Radiology, University of California at San Diego, La Jolla, CA 92093, USA; Epilepsy Center, Cleveland Clinic, Cleveland, OH 44106, USA

## Abstract

Human anterior and posterior hippocampus (aHC, pHC) differ in connectivity and behavioral correlates. Here we report physiological differences. During NREM sleep, the human hippocampus generates sharpwave-ripples (SWR) similar to those which in rodents mark memory replay. We show that while pHC generates SWR, it also generates about as many spindle-ripples (SSR: ripples phase-locked to local spindles). In contrast, SSR are rare in aHC. Like SWR, SSR often co-occur with neocortical theta bursts (TB), downstates (DS), spindles (SS) and upstates (US), which coordinate cortico-hippocampal interactions and facilitate consolidation in rodents. SWR co-occur with these waves in widespread cortical areas, especially fronto-central. These waves typically occur in the sequence TB-DS-SS-US, with SWR usually occurring prior to SS-US. In contrast, SSR occur ∼350 ms later, with a strong preference for co-occurrence with posterior-parietal SS. pHC-SS were strongly phase-locked with parietal-SS, and pHC-SSR were phase-coupled with pHC-SS and parietal-SS. Human SWR (and associated replay events, if any) are separated by ∼5 s on average, whereas ripples on successive SSR peaks are separated by only ∼80 ms. These distinctive physiological properties of pHC-SSR enable an alternative mechanism for hippocampal engagement with neocortex.

**Significance Statement:** Rodent hippocampal neurons replay waking events during sharpwave-ripples in NREM sleep, facilitating memory transfer to a permanent cortical store. We show that human anterior hippocampus also produces sharpwave-ripples, but spindle-ripples predominate in posterior. Whereas sharpwave-ripples typically occur as cortex emerges from inactivity, spindle-ripples typically occur at peak cortical activity. Furthermore, posterior hippocampal spindle-ripples are tightly coupled to posterior parietal locations activated by conscious recollection. Finally, multiple spindle-ripples can recur within a second, whereas sharpwave-ripples are separated by about 5s. The human posterior hippocampus is considered homologous to rodent dorsal hippocampus, which is thought to be specialized for consolidation of specific memory details. We speculate that these distinct physiological characteristics of posterior hippocampal spindle-ripples may support a related function in humans.

## Introduction

Hippocampal sharpwave-ripples (HC-SWR) are striking complexes of local field potentials (LFP) recorded during NREM sleep and rest periods (Buzsáki, 2015). Pyramidal cell firing during HC-SWR tends to reproduce the spatio-temporal patterns established during the preceding active periods; this “replay” is hypothesized to guide cortical activity as it consolidates memories (Wilson and McNaughton, 1994; Skaggs and McNaughton, 1996; Nádasdy et al., 1999). Disrupting HC-SWR impairs memory consolidation (Girardeau et al., 2009; Maingret et al., 2016). The coordination of HC ripples with cortical consolidation is thought to be orchestrated by neocortical LFP graphoelements (NC-GE) such as sleep spindles (SS), down- and upstates (DS/US), and possibly theta bursts (TB) (Gonzalez et al., 2018). Specifically, in rodents, evidence suggests that coordination between HC-SWR and NC-SS (Siapas and Wilson, 1998; Sirota et al., 2003; Peyrache et al., 2009) and DS/US (Battaglia et al., 2004; Mölle et al., 2006; Headley et al., 2016) is central to replay (Diekelmann and Born, 2010; O’Neill et al., 2010; Girardeau and Zugaro, 2011), and recent human data also points to NC-TB, NC-SS, and NC-US coordination being involved in targeted memory reactivation (Göldi et al., 2019). The NC-GE themselves have systematic temporal relations, typically occurring in the order TB→DS→SS→US (Mak-McCully et al., 2017; Gonzalez et al., 2018).

HC-SWR have been extensively studied in rodents. However, two macaque studies have confirmed the presence of HC-SWR during NREM sleep with the typical biphasic negative-positive waves in radiatum, and ∼100 Hz ripples in pyramidale (Skaggs et al., 2007; Ramirez-Villegas et al., 2015). HC-SWR have also been observed in recordings from hippocampal electrodes implanted in humans to identify epileptogenic tissue. In a series of studies, Bragin described HC-SWR but mainly focused on the use of ripples to identify epileptogenic tissue (Bragin et al., 1997, 1999). Axmacher *et al*. (2008) described HC ripples which occurred mainly in waking but were phase-locked to delta. Previously, we described typical SWR in human HC (Jiang et al., submitted-1).

Recently, Staresina et al. (2015) reported that HC ripples in humans during NREM are tightly phase-coupled to hippocampal SS (HC-SS), while presenting no clear evidence for sharp-wave ripples. In this study, we aimed to confirm the presence of ripples associated with HC-SS (henceforth referred to as HC-SSR), and to compare their characteristics and NC-GE associations (if any) with those of HC-SWR, which we reported elsewhere (Jiang et al., submitted-1). Furthermore, since Staresina *et al*. recorded HC-SSR mainly in posterior HC (pHC), whereas previous human HC recordings were mainly in anterior HC (aHC) where most of the human hippocampal volume lies (Destrieux et al., 2013), we examined if HC-SSR selectively occurred in pHC, with HC-SWR dominant in aHC. The possibility that aHC versus pHC have distinct LFP contexts is also suggested by their distinct anatomy and functional correlates (Ranganath and Ritchey, 2012; Poppenk et al., 2013; Strange et al., 2014). Furthermore, since cortical SS preferentially occur in NREM stage N2 compared to N3 (Mak-McCully et al., 2017), we contrasted these stages when performing our NC-GE to HC-GE association analyses. Finally, provided that HC-SSR can be identified, we sought to determine their temporal relationship to HC-SWR, as well as NC-GE.

We report here that, indeed, both SWR and SSR exist in the HC, but with distinct anatomy, cortical relations, and timing. Whereas HC-SSR and HC-SS are largely confined to pHC, HC-SWR predominate in aHC. pHC-SS and pHC-SSR tend to co-occur with NC-SS, especially in parietal sites, while aHC-SWR co-occur with more extensive cortical locations, especially frontal. While aHC-SWR usually occur before SS/US, pHC-SS/SSR tend to occur during NC-SS/US, following NC-TB/DS. Hippocampal and parietal spindles become highly synchronous in about 2 cycles, and both modulate HC ripples. Thus, pHC-SSR interact in close synchrony with parietal NC-SS within the context of more widespread cortical rhythms during NREM. The contrasting anatomy and physiology of pHC-SSR and aHC-SWR provides another possible mechanism for hippocampo-cortical coordination during NREM sleep.

## Methods

### Patient selection

20 patients with long-standing drug-resistant partial seizures underwent SEEG depth electrode implantation in order to localize seizure onset and thus direct surgical treatment (see Table 1 for demographic and clinical information;). Patients were selected from a group of 54 for minimal pathology. The 20 remaining patients included 7 males, aged 29.8±11.9 years old (range 16-58). Electrode targets and implantation durations were chosen entirely on clinical grounds (Gonzalez-Martinez et al., 2013). All patients gave fully informed consent for data usage as monitored by the local Institutional Review Board, in accordance with clinical guidelines and regulations at Cleveland Clinic.

**Table 1.** List of patients, age, sex, handedness, language dominance (LD), NC channel counts, and the lengths sleep period recordings used. Pt: patient. L: Left. R: Right. Dur.: duration. Std dev: standard deviation. T: Total. reused with permission from Jiang et al. (Jiang et al., submitted-1).

| Pt# | Age | Sex | Handedness | # Cortical Channels | HC Seizure-free? | HC Interictal-free? | # Sleep Periods | mean duration of NREM (h) | Std dev NREM (h) | Total N2 Dur. (h) | Total N3 Dur. (h) |
| --- | --- | --- | --- | --- | --- | --- | --- | --- | --- | --- | --- |
| 1 | 20 | M | R | 19 | Y | N | 3 | 2.2 | 0.78 | 2.8 | 3.3 |
| 2 | 51 | F | R | 12 | Y | N | 4 | 7.5 | 1.87 | 27.3 | 0 |
| 3 | 58 | F | R | 24 | Y | N | 4 | 7.1 | 2.44 | 23.1 | 0.30 |
| 4 | 42 | M | L | 17 | Y | N | 4 | 3.1 | 0.33 | 1.8 | 9.0 |
| 5 | 18 | F | L | 21 | Y | Y | 1 | 3.7 |  | 1.9 | 0.9 |
| 6 | 20 | F | R | 21 | Y | N | 3 | 2.7 | 1.49 | 2.6 | 3.0 |
| 7 | 22 | M | LR | 19 | Y | Y | 3 | 3.8 | 1.17 | 2.4 | 3.8 |
| 8 | 30 | F | R | 13 | Y | N | 5 | 5.2 | 1.26 | 6.5 | 14 |
| 9 | 43 | F | R | 12 | Y | N | 4 | 3.1 | 0.45 | 3.7 | 4.4 |
| 10 | 16 | M | R | 17 | Y | Y | 5 | 3.8 | 1.11 | 7.5 | 8.8 |
| 11 | 32 | F | R | 30 | Y | N | 3 | 5.1 | 3.43 | 8.4 | 2.8 |
| 12 | 36 | M | L | 25 | Y | Y | 4 | 5.2 | 1.48 | 11.6 | 6.5 |
| 13 | 21 | F | L | 14 | Y | N | 3 | 6 | 0.47 | 14.7 | 1.1 |
| 14 | 21 | F | R | 14 | Y | N | 8 | 3.7 | 1.1 | 16.9 | 9.3 |
| 15 | 29 | F | R | 17 | Y | N | 4 | 2.5 | 1.31 | 5.4 | 2.7 |
| 16 | 41 | F | R | 19 | Y | N | 3 | 4.6 | 0.72 | 7.4 | 4.4 |
| 17 | 24 | M | R | 21 | Y | N | 3 | 5.1 | 1.88 | 8.9 | 2.9 |
| 18 | 31 | F | R | 15 | Y | N | 6 | 6.1 | 1.18 | 24.2 | 3.8 |
| 19 | 21 | M | R | 15 | Y | Y | 4 | 2.8 | 0.72 | 5.4 | 5.8 |
| 20 | 19 | F | R | 21 | Y | Y | 3 | 4.9 | 2.53 | 5.2 | 6.0 |
| mean | 30 |  |  | 18 | T:20 | T:8 | 4 | 4.4 |  | 9.39 | 4.64 |

### Electrode localization

After implantation, electrodes were located by aligning post-implant CT to preoperative 3D T1-weighted structural MRI with ∼1mm^3^ voxel size (Dykstra et al., 2012), using 3D Slicer (RRID:SCR_005619). This allows visualization of individual contacts with respect to HC cross-sectional anatomy, which was interpreted in reference to the atlas of Duvernoy (Duvernoy, 1988). The assignment of depth contacts to anterior or posterior hippocampus (aHC/pHC) was made with the posterior limit of the uncal head as boundary (Poppenk et al., 2013; Ding and Van Hoesen, 2015). Recordings were obtained from 32 HC contacts, 20 anterior (11 left) and 12 posterior (7 left). In 4 patients, HC recordings were bilateral (3 anterior and 1 posterior), and in 8 patients, ipsilateral anterior and posterior HC were both recorded. The distance of each hippocampal contact from the anterior limit of the hippocampal head was obtained in Freesurfer (RRID:SCR_001847). The CT-visible cortical contacts were then identified as previously described (Jiang et al., submitted-1), in order to ensure that activity recorded by bipolar transcortical pairs is locally generated (Mak-McCully et al., 2015). Electrode contacts were rejected from analysis if they were involved in the early stages of the seizure discharge, had frequent interictal activity or abnormal spontaneous local field potentials. From the total of 2844 contacts implanted in the 20 patients, 366 transcortical pairs (18.3±4.7 per patient) were accepted for further analysis. Polarity of the pairs was adjusted if necessary to “pial surface minus white matter” according to MRI localization, confirmed with decreased high gamma power during surface-negative downstates (see below).

Freesurfer (Dale et al., 1999; Fischl et al., 1999a, 2004) was used to reconstruct from individual MRI scans the cortical pial and inflated surfaces, as well as automatic parcellation of the cortical surface into anatomical areas (Desikan et al., 2006), after a sulcal-gyral alignment process. An average surface for all 20 patients was then generated to serve as the basis of all 3D maps. While each cortical SEEG electrode contact’s location was obtained through direction correlation of CT and MRI as described earlier in this section, we obtained the cortical parcellation labels corresponding to each contact by morphing the Right-Anterior-Superior-oriented anatomical coordinates from individual surfaces to the average surface space (Fischl et al., 1999b). In addition, the nearest vertices on the average surface to the morphed coordinates would be identified for subsequent plotting of 2D projections onto the left lateral view (Fig. 4). For the 2D projections only, to optimize visualization (i.e. minimize multiple contact overlap) while preserving anatomical fidelity, positions of contacts with significant HC-NC GE correlation were allowed to shift within a 5 mm radius. All visualizations were created with custom scripts in MATLAB 2016b.

### Data collection and preprocessing

Continuous recordings from SEEG depth electrodes were made with cable telemetry system (JE-120 amplifier with 128 or 256 channels, 0.016-3000 Hz bandpass, Neurofax EEG-1200, Nihon Kohden) across multiple nights (Table 1) over the course of clinical monitoring for spontaneous seizures, with 1000 Hz sampling rate. The total NREM sleep durations vary across patients; while some difference is expected given intrinsic variability of normal human sleep duration (Carskadon and Dement, 2010) and sleep deprivation in clinical environment, we also confirmed that the percentages of NREM in total sleep from 28 sleeps across 16 of our patients were comparable to (i.e. within 2 standard deviation of) normative data (Moraes et al., 2014) in terms of N2 and N3 durations, with no significant difference in graphoelement occurrence rates between normative and other sleeps (Jiang et al., submitted-1). Recordings were anonymized and converted into the European Data Format (EDF). Subsequent data preprocessing was performed in MATLAB (RRID:SCR_001622); the Fieldtrip toolbox (Oostenveld et al., 2011) was used for bandpass filters, line noise removal, and visual inspection. Separation of patient NREM sleep/wake states from intracranial LFP alone was achieved by previously described methods utilizing clustering of first principal components of delta-to-spindle and delta-to-gamma power ratios across multiple LFP-derived signal vectors (Gervasoni et al., 2004; Jiang et al., 2017), with the addition that separation of N2 and N3 was empirically determined by the proportion of down-states that are also part of slow oscillations (at least 50% for N3 (Silber et al., 2007)), since isolated down-states in the form of K-complexes are predominantly found in stage 2 sleep (Cash et al., 2009).

### Hippocampal graphoelement selection

To identify hippocampal spindles, we applied a previously reported spindle detector (Mak-McCully et al., 2017) to hippocampal LFP signals from NREM: a 10–16 Hz bandpass (zero-phase shift filter with transition bands equal to 30% of the cutoff frequencies) was applied to the LFP data, and the analytic amplitude of the bandpassed signal was convolved with an average Tukey window of 600 ms. A channel-wise cutoff set at mean +2 s.d. was applied to identify local maxima that would be “peaks” of putative spindles, with spindle edges defined at 50 % of peak amplitude. The putative HC-SS segments were accepted for subsequent analyses if they were longer than 400 ms, had less than 14 dB power in the nearby frequency bands (4-8 Hz and 18-30 Hz), and contained at least 3 unique oscillation peaks with 40-100 ms gaps.

Hippocampal SWR were identified by starting with the top 20% of peaks in root-mean-square amplitude of 60-120 Hz LFP over a moving 20 ms window, then requiring that putative SWR events: (1) be >200 ms apart; (2) have >3 distinct ripple peaks in the HC LFP signal (low-passed at 120 Hz); and (3) be associated with an LFP matching hand-marked sharp-waves in the same subject and channel (Jiang et al., submitted-1). RMS peaks potentially arising from artifacts or epileptiform activity were rejected by detecting sharp LFP transients and setting a threshold that separated hand-marked HC-SWR from interictal events (Jiang et al., submitted-1). Prior to the SWR similarity evaluation, putative ripples were also examined for whether a given ripple center falls within a hippocampal spindle duration in the same signal; if positive, such ripples were then deemed SSR. Since a single ripple could qualify as both SWR and SSR, such a ripple would be denoted as SXR where appropriate.

### Cortical graphoelement selection

Automatic cortical theta burst (TB) detection was performed as previously described (Gonzalez et al., 2018): a 5-9 Hz zero-phase shift bandpass filter was applied to the LFP data, and a channel-wise cutoff set at mean +3 s.d. was applied to each channel’s Hilbert envelope (smoothed with 300 ms Gaussian kernel) to identify local maxima that would be “peaks” of putative TBs, and make start/stop edge cut-offs at mean +1 s.d.. The resulting TB were only accepted for further analyses if their durations fall between 400 ms and 1000 ms, with their number-of-zero-crossings-based frequency estimate falling within the bandpass frequency range as well.

Automatic cortical spindle (SS) detection was performed in the same matter as previously described for hippocampal spindles. Downstates (DS) and upstates (US) were identified as follows: for each cortical LFP signal in NREM, a zero-phase filter from 0.1 to 4 Hz was applied. Consecutive zero crossings of opposite slope separated by 250 to 3000 ms were then selected as delineating putative graphoelements. For each putative graphoelement, the amplitude peak between zero crossing was computed; only the top 10% of peaks were retained. The polarity of each signal was inverted if necessary to assure that DS were negative, as confirmed with a high gamma (70-190 Hz) power decrease exceeding 1 dB within ±250 ms of the negative DS peaks.

### Experimental design and statistical analysis

Statistical tests that involve multiple comparisons in this and subsequent Methods subsections all have α = 0.05 post-FDR correction (Benjamini and Hochberg, 1995). The FDR correction procedure was implemented as follows: for a total of N hypotheses, each with corresponding p-values P_*i*_ (*i* ∈ N), which are sorted in ascending order to identify P_*k*_, *k* being the largest *i* for which P_*i*_ ≤ (*i* / N) * α. All hypothesis with p-values less than or equal to P*k* would then be rejected. The striking positive skewness both in general and in specific categories of the NC-GE histograms (Fig. 3-1) suggests high variability in co-occurrence rates of NC-GE with HC-SS and with HC-SSR. Therefore, to account for sampling error, we created linear mixed effect (LME) models for several analyses as described below.

We compared the occurrence rates of HC-SSR/SS from different NREM stages (N2 vs. N3) and from different sources (aHC vs. pHC) with both paired t-tests and LME models to account for patient-wise variability. For LME, our base models have either HC-SSR or HC-SS rates as the response variable, with AP (aHC/pHC) and Stage (N2/N3) as categorical predictor variables. The full model includes interaction between predictor variables and all possible random slope/intercept terms (GE Rate ∼ 1 + AP*Stage + (AP | Patient ID) + (Stage | Patient ID) + (1 | Patient ID)), with reduced models’ fits being compared to the full model fit via likelihood ratio tests. The best-fit models for both HC-SSR and HC-SS turned out to be the following: HC-GE Rate ∼ 1 + AP*Stage + (AP | Patient ID) + (1 | Patient ID).

Similarly, we compared the occurrence rates of HC-SWR with those of HC-SSR in different NREM stages and from different hippocampal origins, while accounting for random effect due to patient identity. Our full LME model has HC-GE rates as the response variable, with three categorical predictor variables and all possible random slopes, plus random intercept due to patient identity. By stepwise removal of individual terms from the full model and comparing each reduced model with the one prior to it via likelihood ratio tests, we arrived at the best fit model of GE Rate ∼ 1 + GE Type*AP*Stage + (GE Type | Patient ID).

We also evaluated whether a significant difference between SSR/SWR ratio in the first versus second half of NREM sleep could be found, while accounting for variance due to patient identity. Our base LME model has SSR/SWR ratio as the response variable, and “sleep halves” as the categorical predictor variable. After comparing models with random intercept and/or slope with base model, we found the best fit to be the base model (Ratio ∼ 1 + SleepHalves).

For time-frequency analysis in Figure 1, spectral content of the LFP data from hippocampal channels were evaluated using EEGLAB (RRID:SCR_007292) routines that applied wavelet transforms (Delorme and Makeig, 2004). Spectral power was calculated over 1 to 200 Hz across each individual time period (“trial”) centered around HC-SWR ripple centers, HC-SSR ripple centers, or HC-SS onsets found in NREM by convoluting each trial’s signal with complex Morlet wavelets, and averages were then taken across trials. The resulting time-frequency matrices were normalized with respect to the mean power at each chosen frequency and masked with two-tailed bootstrap significance at α = 0.05, with the pre-trigger times (−2000 ms to −1500 ms) as baseline.

**Figure 1.**
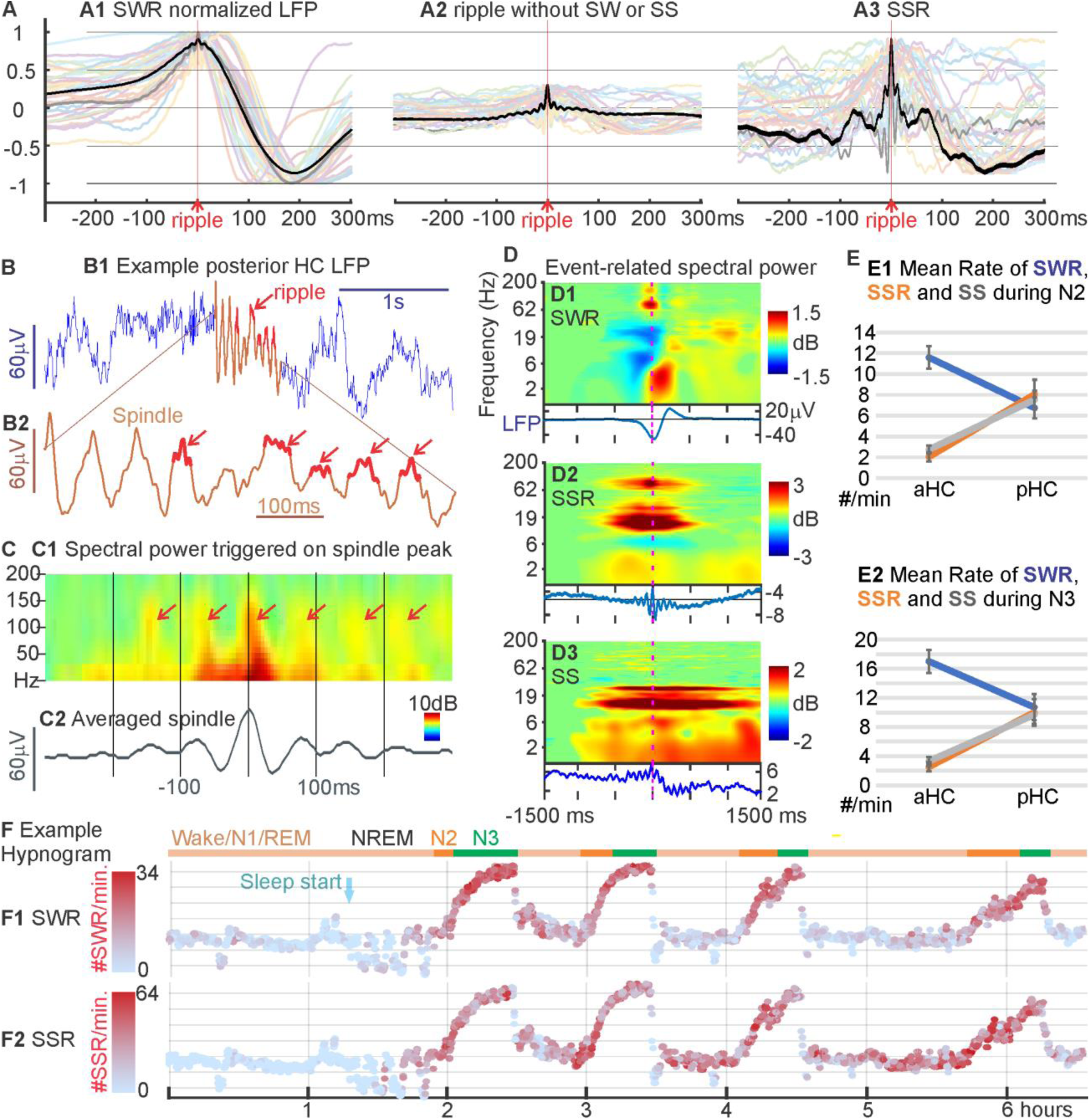
Hippocampal spindles (HC-SS) co-ordinate ripples and are found predominantly in posterior HC. ***A***. Overlaid average waveforms of HC-SWR (***A1***), of ripples not coupled to sharpwaves (***A2***), and of HC-SSR (***A3***) across hippocampal contacts from 32 different patients, with one representative patient bolded in black for clarity. The non-normalized, across-patient mean LFP peak-to-peak amplitude of the sharp wave (black trace in ***A***) is 127.1 μV, much greater than the corresponding peak amplitudes of 8.284 and 14.79 μV in ***B*** and ***C***, respectively. ***B***. Example LFP trace of a ∼14 Hz HC-SS (red section) from bipolar recording in the posterior HC (***B1***), with expanded time-base plot of the spindle shown in ***B2***; 80-90 Hz ripples occur preferentially at some peaks of the spindle waves. ***C***. Example averaged time-frequency plot triggered on the center peak of HC-SS from the same channel in ***B*** (***C1***, n=1086) and the corresponding average LFP (***C2***). ***D***. Event-related spectral power is contrasted for SWR (***D1***), SSR (***D2***), and SS (***D3***) from the same HC site, with corresponding average LFP traces below the time-frequency plots. The high frequency ripple burst is seen for SWR and SSR but not prominently so in SS. Prolonged spindle activity is seen for SSR and SS but not SWR. ***E***. HC-SWR preferentially occur in aHC while HC-SS/SSR prefer pHC. Each panel marks mean rates of HC-SWR/SSR/SS occurrence across patients in N2 (***E1***) or N3 (***E2***), for aHC and pHC. ***F***. Example state plots showing the separation of NREM sleep stages N2 and N3 from waking/N1/REM in ∼6.5-hour LFP recording, using the first principal component derived from vectors (one per cortical bipolar SEEG channel) of frequency power ratios (0.5-3 Hz over 0.5-16 Hz) (Gervasoni et al., 2004; Jiang et al., 2017). SWR rate (***F1***) and SSR rate (***F2***) are color coded with red intensity, and N2/N3 periods are marked with orange/green horizontal lines, respectively. Light blue arrow marks the beginning of NREM sleep. Similar illustration for HC-SS rate over time can be found in Extended Data Figure 1-1. Error bars in ***E*** denote standard errors of the mean.

Peri-stimulus time histograms were constructed for each HC-NC channel pair, separately for each graphoelement (GE; TB, SS, DS, and US), and for each sleep-stage (N2 or N3), each histogram comprising the occurrence times of a given NC-GE during the ±2 s interval surrounding midpoints of HC-SSR and onsets of HC-SS (see Fig. 3 for examples). Altogether, there were 598 unique HC-NC pairs (458 ipsilateral, 140 contralateral), and 16 histograms for each pair (one for each sleep stage, times each of the 4 GE types, for HC-SS or SSR), for a total of 9568 histograms. On average, each of these histograms plots the occurrence times of 1386±2565 NC-GE (+3.86 skewness) with respect to 861±1581 HC-SSR (+3.57 skewness) and to 1022±1642 HC-SS (+3.61 skewness). Overall a total of 13.2 million NC-GE events, 4.12 million HC-SSR, and 4.89 million HC-SS were plotted.

The significance of peaks and troughs in each histogram was tested by comparing them to distributions derived from histograms constructed under the null hypothesis of no relationship between the NC-GE and HC-SS/SSR using the following randomization procedure. Null-hypothesis histograms (N=1000) were constructed of NC-GE occurrences relative to a number of random times equal to the number of HC-SS/SSR (examples shown in Fig. 3). For each 200 ms time bin with 100 ms overlap comprising the 4-second trial duration, the actual counts are then compared to the distribution under the null hypothesis, followed by FDR correction. Similar peri-stimulus histograms and randomization tests were also constructed with aHC-SWR as triggers and pHC-SWR/SSR as tallied events (Fig. 2*A*).

**Figure 2.**
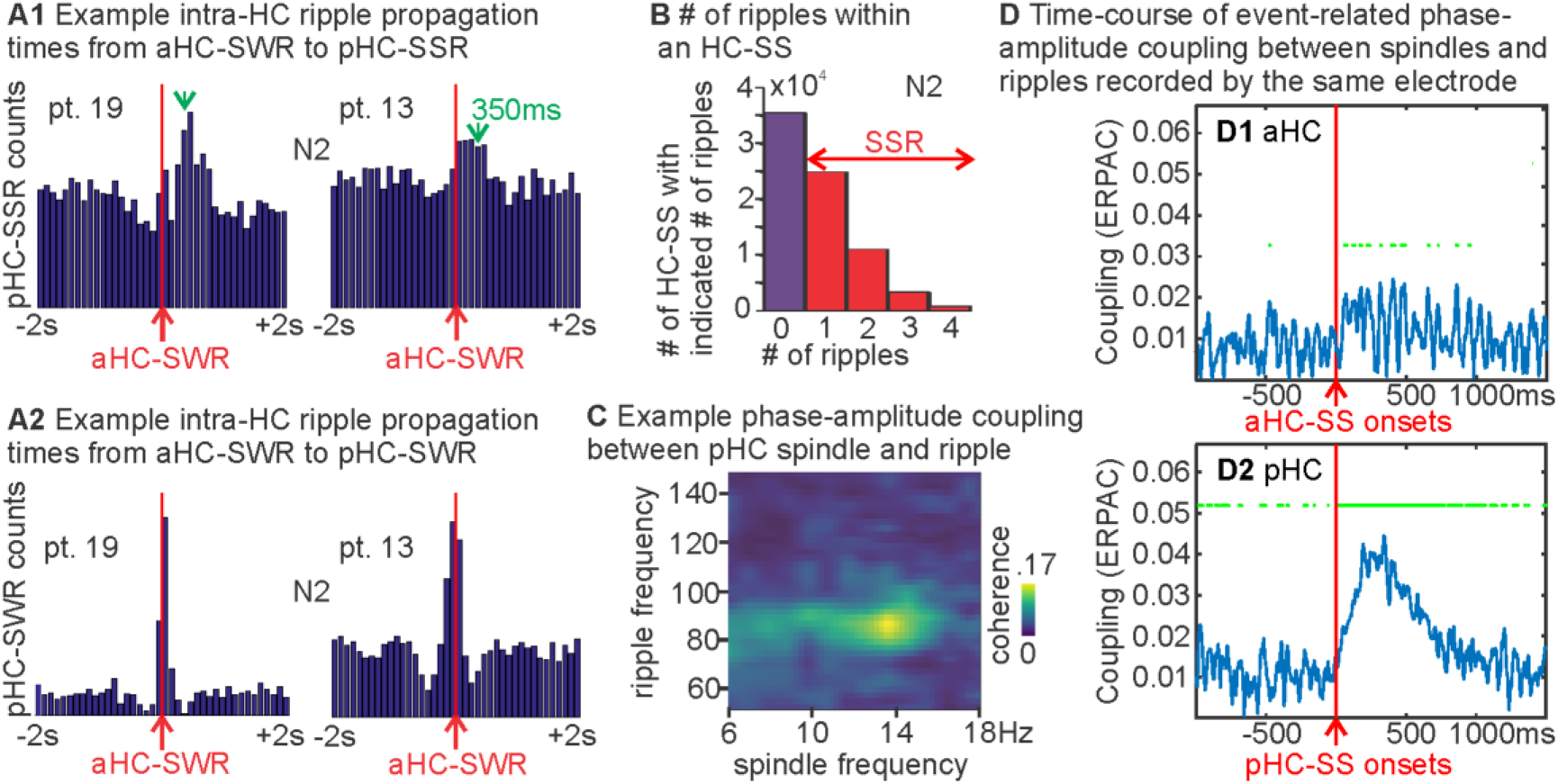
Temporal relations of ripples and spindles, within and between hippocampal locations. ***A***. Peri-stimulus time histograms with aHC-SWR as triggers and either pHC-SSR (***A1***) or pHC-SWR (***A2***) as tallied events, with HC-SXR excluded from ***A1***; in N2, pHC-SSR tend to follow aHC-SWR by ∼350 ms (marked with green arrows in ***A***), and aHC/pHC-SWR tend to co-occur within ±200 ms. Similar results for N3 can be found in Extended Data Fig. 2-1*F-G*. ***B***. Histograms of ripple counts across all HC-SS in N2 (N3 results are in Fig. 2-1*H*). Not shown are the HC-SS with ≥5 ripples (n=256 in N2, max 9). ***C***. Phase-amplitude coupling within HC between HC-SS phase and ripple amplitude (bandpassed between 50 and 150 Hz). ***D***. The time-course of intra-hippocampal event-related phase-amplitude coupling (ERPAC) between spindle (10-16 Hz) phase and ripple (60-120 Hz) amplitude, computed over all HC-SS in NREM across all patients, separately for aHC (D1) and pHC (D2). Events are aligned at HC-SS starts (red vertical lines). Green line segments mark the time segments with significant coupling (p < 0.001). The same pattern is observed if pHC events are randomly sub-selected to have the same number of events involved as aHC (Fig. 2-1*E*).

**Figure 3.**
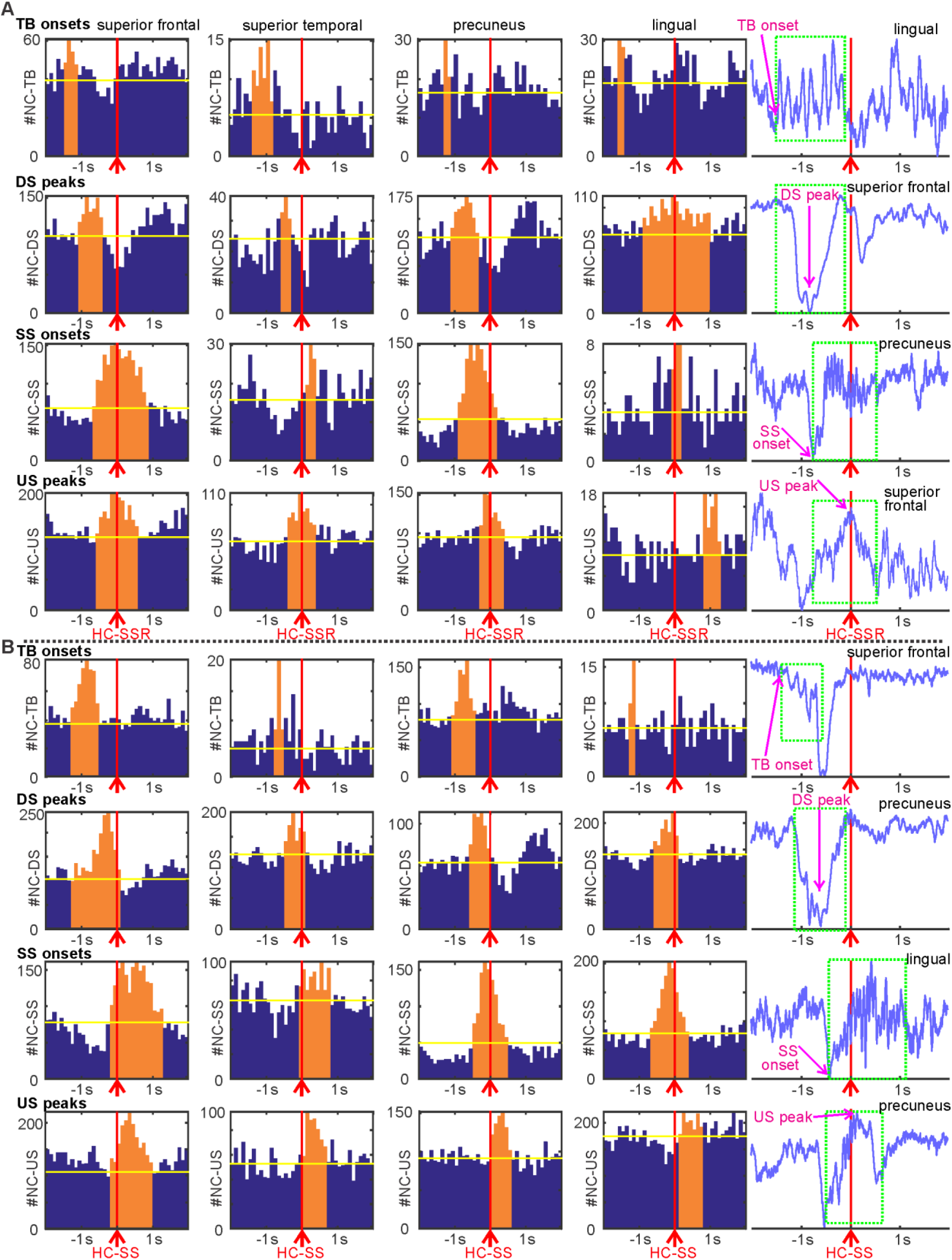
Neocortical graphoelements (NC-GE) in relation to HC-SSR (***A***) and HC-SS (***B***), for example channel-pairs. Each row of histograms is for a different type of NC-GE: theta burst onsets (TB), downstate peaks (DS), spindle onsets (SS), and upstate peaks (US), with the rightmost panel showing an example LFP trace containing the corresponding NC-GE (marked in green dash line boxes). Each column of plots shows example histograms with significant temporal correlations between HC-SS/SSR and NC-GE from the same NC region in (from left to right) frontal, temporal, parietal, and occipital lobes. Magenta vertical lines indicate trigger (HC-SS/SSR) location for the peri-stimulus histograms. Orange bars indicate the time ranges with both peak NC-GE occurrence rate and significant correlation. Yellow horizontal bars mark the average expected NC-GE counts from randomized control distributions. A summary of histogram contents can be found in Extended Data Figure 3-1.

The latencies of the largest significant peaks identified in the individual histograms constructed as described above, were used to create summary histograms for each GE, sleep stage, and HC-SS/SSR origin in anterior versus posterior HC. These are plotted in Fig. 4, and tabulated in Table 3. Each of the 16 histograms of histogram peaks (N2/N3 × aHC/pHC × 4 NC-GE) for HC-SSR summarizes the significant latencies from 78±41 HC-NC channel pair histograms, comprising a total of 1.37 × 10^5^±1.45 × 10^5^ HC-NC events (range 7.18 × 10^3^-4.04 × 10^5^). Each of the 16 histograms of histogram peaks for HC-SS summarizes the significant latencies from 126±63 HC-NC channel pair histograms, comprising a total of 2.46 × 10^5^±2.07 × 10^5^ HC-NC events (range 1.24 × 10^4^-6.23 × 10^5^). To test if, overall, a given type of cortical graphoelement significantly precedes or follows HC-SS/SSR, two-tailed binomial tests with chance probability of 0.5 were performed on the number of channel-pairs with peak latencies in the 2000 ms before vs the 2000 ms after the reference HC-SS/SSR. Since certain graphoelements, such as cortical theta, tend to yield peak latencies centered around zero, we also tested with Kolmogorov-Smirnov (KS) tests whether, overall, for a given NC-GE, the distribution of peak latencies significantly related to HC-SS/SSR differs from chance.

**Table 2.** Hippocampal SEEG electrode coverage and occurrence rates of hippocampal graphoelements. SD: standard deviation.

| Pt# | L aHC | L pHC | R aHC | R pHC | SWR Rate, N2 aHC (/min) | SWR Rate, N2 pHC (/min) | SWR Rate, N3 aHC (/min) | SWR Rate, N3 pHC (/min) | SSR Rate, N2 aHC (/min) | SSR Rate, N2, pHC (/min) | SSR Rate, N3, aHC (/min) | SSR Rate, N3, pHC (/min) | HC-SS Rate, N2, aHC (/min) | HC-SS Rate, N2, pHC (/min) | HC-SS Rate, N3, aHC (/min) | HC-SS Rate, N3, pHC (/min) |
| --- | --- | --- | --- | --- | --- | --- | --- | --- | --- | --- | --- | --- | --- | --- | --- | --- |
| 1 | x |  |  |  | 11 |  | 19 |  | 6.3 |  | 9.4 |  | 5.8 |  | 9 |  |
| 2 | x |  |  |  | 1.6 |  |  |  | 0.67 |  |  |  | 1.7 |  |  |  |
| 3 | x | x |  |  | 11 | 12 | 24 | 25 | 1.4 | 3.4 | 1.4 | 5.6 | 2.8 | 5.4 | 2.6 | 9.3 |
| 4 | x | x | x |  | 10 | 4.3 | 11 | 4.3 | 1.4 | 3.8 | 1.1 | 1.6 | 2.6 | 5.5 | 2.4 | 3.2 |
| 5 |  |  | x | x | 21 | 3.7 | 38 | 14 | 1.3 | 7 | 1.9 | 9.9 | 1.4 | 4 | 1.8 | 7.2 |
| 6 |  |  | x | x | 9 | 12 | 15 | 17 | 4.6 | 18 | 5.2 | 22 | 6.1 | 10 | 7.6 | 14 |
| 7 | x |  |  |  | 11 |  | 13 |  | 0.27 |  | 0.26 |  | 0.52 |  | 0.43 |  |
| 8 |  |  | x |  | 19 |  | 20 |  | 1.2 |  | 1.3 |  | 1.4 |  | 1.9 |  |
| 9 | x | x |  |  | 13 | 7.9 | 16 | 7.2 | 1.1 | 7.2 | 1.5 | 11 | 2.2 | 5.9 | 2.8 | 9.3 |
| 10 |  |  | x |  | 13 |  | 18 |  | 2.7 |  | 2.2 |  | 2.7 |  | 2.7 |  |
| 11 | x |  | x | x | 7.3 | 7.7 | 12 | 12 | 0.44 | 1.1 | 0.49 | 1.5 | 1.5 | 2.4 | 1.1 | 2.8 |
| 12 |  |  |  | x |  | 5.5 |  | 7 |  | 13.4 |  | 19 |  | 12 |  | 17 |
| 13 | x | x | x |  | 13 | 6.2 | 19 | 4.9 | 2.6 | 8 | 2.4 | 8.2 | 3.6 | 9.8 | 3 | 13 |
| 14 |  | x |  |  |  | 11 |  | 15 |  | 7 |  | 7.9 |  | 11 |  | 12 |
| 15 |  | x |  | x |  | 3.6 |  | 7.2 |  | 7.1 |  | 8.6 |  | 7.2 |  | 7.6 |
| 16 |  |  | x |  | 9.4 |  | 11 |  | 1.6 |  | 2.2 |  | 2.2 |  | 3 |  |
| 17 | x |  |  |  | 23 |  | 24 |  | 0.75 |  | 0.89 |  | 1.4 |  | 1.4 |  |
| 18 | x |  |  |  | 7.8 |  | 10 |  | 1.2 |  | 1.1 |  | 1.9 |  | 1.9 |  |
| 19 | x | x |  |  | 12 | 2.6 | 19 | 6.1 | 7.7 | 14 | 9.7 | 16 | 6.6 | 9.7 | 9.3 | 12 |
| 20 |  |  | x |  | 11 |  | 12 |  | 1.3 |  | 3 |  | 2.9 |  | 5.6 |  |
| mean |  |  |  |  | 12 | 7 | 18 | 11 | 2.1 | 8.2 | 2.8 | 10 | 2.8 | 7.5 | 3.5 | 9.8 |
| SD |  |  |  |  | 1.1 | 1.0 | 1.6 | 1.8 | 0.44 | 1.4 | 0.62 | 1.9 | 0.36 | 0.87 | 0.59 | 1.3 |

**Table 3.** Relation of NC-GE to HC-SSR and HC-SS occurrence times. Separate values and statistical significance tests are shown for NREM sleep stages N2 vs. N3, for anterior vs. posterior HC, and for different NC graphoelements (TB-theta bursts; SS-sleep spindles; DS-downstates; US-upstates). Tests indicate if there was a significant association between the times of occurrence of the HC-SSR/HC-SS and GE (fourth and eighth columns); if the NC-GE occurred significantly before the HC-SSR/HC-SS (negative numbers) or before, when measuring the peak latency as the mean value across cortico-HC pairs (fifth and ninth columns), or as the peak of an extreme value distribution fitted over the ±2000 ms histogram-of-histograms in Figs. 4*A* and 4-1 (sixth column) or in Figs. 4*B* and 4-2 (tenth column). The seventh and eleventh columns show if there is a significant difference in the number of NC-GE occurring 2000 ms before vs. 2000 ms after the HC-SSR/HC-SS. Extremely small (< 0.0001) p-values are represented by 0 for clarity. While NC-DS was significantly earlier than pHC-SSR in N3, but not in N2, it should be noted that NC-DS distribution appeared bimodal for N2 pHC-SSR, which likely distorted the mean and distribution peak estimations: the larger pre-trigger peak of the NC-DS latency distribution for N2 pHC-SSR, like the NC-DS distribution peak in N3, preceded that of the NC-SS latency distribution (∼700 ms pre-SSR for NC-DS vs. 353 ms for NC-SS) (Fig. 4*A*).

| NC-GE Type | Sleep Stage | HC Longitudinal Axis | NC-GE v. HC-SSR related (KS test p-value) | Peak Latency from HC-SSR (ms), Mean | Peak Latency from HC-SSR (ms), DistPeak | Non-zero Latency from HC-SSR, binomial p-value | NC-GE v. HC-SS related (KS test p-value) | Peak Latency from HC-SS (ms), Mean | Peak Latency from HC-SS (ms), DistPeak | Non-zero Latency from HC-SS, binomial p-value |
| --- | --- | --- | --- | --- | --- | --- | --- | --- | --- | --- |
| TB onsets | N2 | anterior | 0 | -398 | -741 | 0 | 0 | -164 | -392 | 0 |
|  |  | posterior | 0.002 | -386 | -1074 | 0.011 | 0 | -455 | -698 | 0 |
|  | N3 | anterior | 0.129 | 123 | 206 | 1 | 0 | -96 | -338 | 0.006 |
|  |  | posterior | 0.654 | -131 | -346 | 0.388 | 0 | -512 | -646 | 0 |
| DS peaks | N2 | anterior | 0 | -185 | -575 | 0 | 0 | 55 | -65 | 0.232 |
|  |  | posterior | 0 | 38 | 31 | 0.519 | 0 | 105 | -167 | 0.001 |
|  | N3 | anterior | 0.216 | -95 | -426 | 0.117 | 0 | 148 | -57 | 1 |
|  |  | posterior | 0 | -288 | -489 | 0.001 | 0 | -64 | -184 | 0 |
| SS onsets | N2 | anterior | 0 | -16 | -163 | 0.213 | 0 | 80 | 73 | 0.086 |
|  |  | posterior | 0 | -211 | -353 | 0 | 0 | -19 | -106 | 0.368 |
|  | N3 | anterior | 0.017 | -32 | -142 | 0.728 | 0.003 | -10 | 12 | 0.377 |
|  |  | posterior | 0 | -241 | -425 | 0 | 0 | -45 | -32 | 0.013 |
| US peaks | N2 | anterior | 0 | 35 | 93 | 0.221 | 0 | 348 | 534 | 0 |
|  |  | posterior | 0 | 68 | 62 | 0.643 | 0 | 165 | 212 | 0 |
|  | N3 | anterior | 0.013 | -131 | -180 | 0.609 | 0 | 338 | 568 | 0 |
|  |  | posterior | 0 | -49 | -71 | 0.699 | 0 | 199 | 111 | 0 |

To evaluate the phase relationship between cortical and hippocampal spindle oscillations, we computed phase-locking values (PLV) (Lachaux et al., 1999) between each cortical and hippocampal channel pairing over 3-second trials centered on all hippocampal spindle event starts in NREM (N2+N3). Only hippocampal graphoelements that overlapped with the cortical spindle for at least 160 ms were included, in order to ensure that the cortical and hippocampal spindles overlap for at least 1 spindle cycle, thus allowing more accurate estimates of phase-locking. We also computed PLV for the same cortical-hippocampal channel pairs over the same number of trials centered on random times in NREM to create a baseline estimate, and for each non-overlapping 50 ms time bin, a two-sample t-test was performed between the actual PLV and the baseline estimate, with the resulting p-values undergoing FDR correction. A given channel pair would be considered significantly phase-locking if: 1, more than 40 trials were used in the PLV computation, since small sample size is known to introduce bias (Aydore et al., 2013); 2, at least 3 consecutive time bins yield post-FDR p-values below 0.05.

To evaluate phase-amplitude coupling between cortical spindle phase and hippocampal ripple amplitude, we adopted a Modulation Index method as follows (Tort et al., 2010): for each HC-NC channel pair with significant SS PLV, we first computed instantaneous phase and amplitude through Hilbert transform for each trial (as previously obtained for PLV analysis), with NC-SS phase angles in 10 degree bins to create a corresponding mean amplitude distribution across trials. For statistical evaluation of observed PAC, we then computed the Modulation Index (i.e. dividing the Kullback-Leibler distance of the observed amplitude distribution from the uniform distribution by log (N), N being the number of trials) for this channel pair, and performed permutation tests by generating 200 additional Modulation Indices with random pairings of the NC-SS phase and HC ripple amplitude (so that phase from one trial is matched to amplitude from another trial). The permutation test results were then FDR-corrected.

For intra-hippocampal coupling between ripple-frequency amplitude (70-100 Hz) and spindle (10-16 Hz) phase, we applied previously published methods for evaluating event-related phase-amplitude coupling (ERPAC) and estimating subsequently ERPAC significance over time from intracranial LFP data (Voytek et al., 2013). The events were NREM hippocampal spindle starts, and each ERPAC trial covers -1000 ms to +1500 ms peri-event. Each trial was also marked as either overlapping with NC-SS (by more than one spindle cycle) or not. Statistical evaluation of significant coupling was done using permutation tests that kept the actual analytic amplitude and phase values at each time point, but randomized the trial labels 1,000 times so that the amplitude values are matched to phase values from random trials. The resulting p-values for each time point then underwent FDR correction. To generate the comodulogram in Fig. 2*C*, we concatenated the hippocampal data across spindle events and applied previously published phase-amplitude coupling algorithm (Colgin et al., 2009) implemented within the Pactools software package (Tour et al., 2017) under Python 3.6 (RRID:SCR_008394).

In order to explore the anatomical distribution of HC-NC GE co-occurrences, we tested the following: 1. whether the distribution of significantly SSR- or HC SS-coupled NC sites was different from chance for a given HC site; 2. whether the significantly-different-from chance distributions of NC sites from 1 would also differ between aHC/pHC site pairs; 3. whether the NC sites showing significant HC-NC correlation with one NC-GE type tended to also show correlation with another NC-GE type.

For 1, we performed a chi-square test of homogeneity for each HC site with regard to each GE, i.e. for a given patient with multiple NC sites, a 2 × N contingency table was made for each NC site, N being the number of NC sites, one row of table containing the number of NC-GE overlapping with (i.e. occurring within 1500 ms of) HC-SS/SSR for each NC, the other containing the number of NC-GE that do not. A significant chi-square would therefore indicate a significant non-random distribution for that particular HC site and NC-GE type.

For 2, we performed Wilcoxon signed rank tests on the data obtained in 1 to evaluate whether for a given NC-GE type (for which 1 yielded significance for both members of an HC site pair), the proportions of GE from significantly coupled NC sites that overlapped with aHC-SSR/SS differ from the proportions for pHC-SSR/SS. Each test required that both members of a HC site pair have a significant chi-square test result from 1.

We further explored the spatial distribution of HC-NC relationship by tallying the proportion of significant HC-NC channel pairs across different NC regions, with respect to different NC-GE types and different HC-SS/SSR sources (Table 4, Table 5). To characterize the apparent variability in Tables 4 and 5, we performed 4-way ANOVAs to compare the main effects of GE type (TB, SS, DS, US), NREM stage (N2 vs. N3), aHC or pHC origin of HC-SS/SSR, and NC ROIs (coverage listed in Table 4-1) as well as the 2-way and 3-way interaction effects (Table 6, Table 7). To evaluate the differences in NC-HC coupling between SWR and SSR, we combined the SSR data with data from the equivalent analysis in our study on SWR (Jiang et al., submitted-1), and performed another 4-way ANOVA with the following factors: SSR/SWR as the participant of NC-HC coupling, NREM stage, aHC/pHC origin of HC-SWR/SSR, and NC ROIs.

**Table 4.**
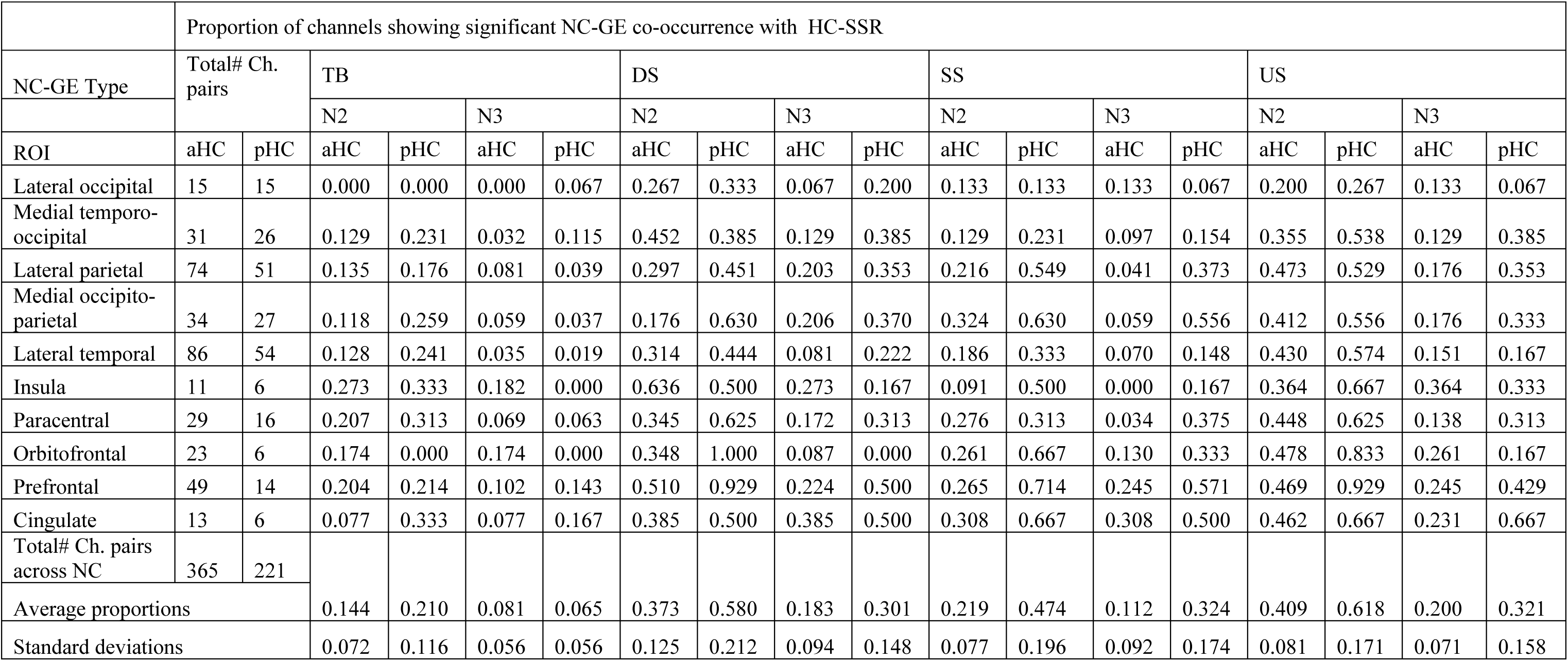
Numbers of HC-NC channel pairs by NC regions of interest (ROIs), and proportions of HC-NC channel pairs with significant NC-GE/HC-SSR relationships across NC ROIs. Ch. pairs: HC-NC channel pairs. Details on Freesurfer parcellations included in each ROI can be found in Extended Data Table 4-1.

|  | Proportion of channels showing significant NC-GE co-occurrence with HC-SSR |  |  |  |  |  |  |  |  |  |  |  |  |  |  |  |  |  |
| --- | --- | --- | --- | --- | --- | --- | --- | --- | --- | --- | --- | --- | --- | --- | --- | --- | --- | --- |
| NC-GE Type | Total# Ch. pairs |  | TB |  |  |  | DS |  |  |  | SS |  |  |  | US |  |  |  |
|  |  |  | N2 |  | N3 |  | N2 |  | N3 |  | N2 |  | N3 |  | N2 |  | N3 |  |
| ROI | aHC | pHC | aHC | pHC | aHC | pHC | aHC | pHC | aHC | pHC | aHC | pHC | aHC | pHC | aHC | pHC | aHC | pHC |
| Lateral occipital | 15 | 15 | 0.000 | 0.000 | 0.000 | 0.067 | 0.267 | 0.333 | 0.067 | 0.200 | 0.133 | 0.133 | 0.133 | 0.067 | 0.200 | 0.267 | 0.133 | 0.067 |
| Medial temporo-occipital | 31 | 26 | 0.129 | 0.231 | 0.032 | 0.115 | 0.452 | 0.385 | 0.129 | 0.385 | 0.129 | 0.231 | 0.097 | 0.154 | 0.355 | 0.538 | 0.129 | 0.385 |
| Lateral parietal | 74 | 51 | 0.135 | 0.176 | 0.081 | 0.039 | 0.297 | 0.451 | 0.203 | 0.353 | 0.216 | 0.549 | 0.041 | 0.373 | 0.473 | 0.529 | 0.176 | 0.353 |
| Medial occipito-parietal | 34 | 27 | 0.118 | 0.259 | 0.059 | 0.037 | 0.176 | 0.630 | 0.206 | 0.370 | 0.324 | 0.630 | 0.059 | 0.556 | 0.412 | 0.556 | 0.176 | 0.333 |
| Lateral temporal | 86 | 54 | 0.128 | 0.241 | 0.035 | 0.019 | 0.314 | 0.444 | 0.081 | 0.222 | 0.186 | 0.333 | 0.070 | 0.148 | 0.430 | 0.574 | 0.151 | 0.167 |
| Insula | 11 | 6 | 0.273 | 0.333 | 0.182 | 0.000 | 0.636 | 0.500 | 0.273 | 0.167 | 0.091 | 0.500 | 0.000 | 0.167 | 0.364 | 0.667 | 0.364 | 0.333 |
| Paracentral | 29 | 16 | 0.207 | 0.313 | 0.069 | 0.063 | 0.345 | 0.625 | 0.172 | 0.313 | 0.276 | 0.313 | 0.034 | 0.375 | 0.448 | 0.625 | 0.138 | 0.313 |
| Orbitofrontal | 23 | 6 | 0.174 | 0.000 | 0.174 | 0.000 | 0.348 | 1.000 | 0.087 | 0.000 | 0.261 | 0.667 | 0.130 | 0.333 | 0.478 | 0.833 | 0.261 | 0.167 |
| Prefrontal | 49 | 14 | 0.204 | 0.214 | 0.102 | 0.143 | 0.510 | 0.929 | 0.224 | 0.500 | 0.265 | 0.714 | 0.245 | 0.571 | 0.469 | 0.929 | 0.245 | 0.429 |
| Cingulate | 13 | 6 | 0.077 | 0.333 | 0.077 | 0.167 | 0.385 | 0.500 | 0.385 | 0.500 | 0.308 | 0.667 | 0.308 | 0.500 | 0.462 | 0.667 | 0.231 | 0.667 |
| Total# Ch. pairs across NC | 365 | 221 |  |  |  |  |  |  |  |  |  |  |  |  |  |  |  |  |
| Average proportions |  |  | 0.144 | 0.210 | 0.081 | 0.065 | 0.373 | 0.580 | 0.183 | 0.301 | 0.219 | 0.474 | 0.112 | 0.324 | 0.409 | 0.618 | 0.200 | 0.321 |
| Standard deviations |  |  | 0.072 | 0.116 | 0.056 | 0.056 | 0.125 | 0.212 | 0.094 | 0.148 | 0.077 | 0.196 | 0.092 | 0.174 | 0.081 | 0.171 | 0.071 | 0.158 |

**Table 5.** Numbers of HC-NC channel pairs by NC regions of interest (ROIs), and proportions of HC-NC channel pairs with significant relationships between NC-GE and HC-SS. Ch. pairs: HC-NC channel pairs.

|  |  |  | Proportion of channels showing significant NC-GE co-occurrence with HC-SS |  |  |  |  |  |  |  |  |  |  |  |  |  |  |  |
| --- | --- | --- | --- | --- | --- | --- | --- | --- | --- | --- | --- | --- | --- | --- | --- | --- | --- | --- |
| NC-GE Type | Total# Ch. pairs |  | TB |  |  |  | DS |  |  |  | SS |  |  |  | US |  |  |  |
|  |  |  | N2 |  | N3 |  | N2 |  | N3 |  | N2 |  | N3 |  | N2 |  | N3 |  |
| ROI | aHC | pHC | aHC | pHC | aHC | pHC | aHC | pHC | aHC | pHC | aHC | pHC | aHC | pHC | aHC | pHC | aHC | pHC |
| Lateral occipital | 15 | 15 | 0.200 | 0.133 | 0.133 | 0.000 | 0.200 | 0.600 | 0.067 | 0.467 | 0.133 | 0.200 | 0.133 | 0.133 | 0.467 | 0.333 | 0.400 | 0.600 |
| Medial temporo-occipital | 31 | 26 | 0.194 | 0.269 | 0.161 | 0.231 | 0.613 | 0.500 | 0.194 | 0.423 | 0.194 | 0.192 | 0.129 | 0.154 | 0.581 | 0.538 | 0.258 | 0.538 |
| Lateral parietal | 74 | 51 | 0.432 | 0.412 | 0.176 | 0.176 | 0.757 | 0.706 | 0.311 | 0.608 | 0.324 | 0.706 | 0.095 | 0.353 | 0.797 | 0.627 | 0.365 | 0.804 |
| Medial occipito-parietal | 34 | 27 | 0.324 | 0.370 | 0.147 | 0.185 | 0.529 | 0.630 | 0.265 | 0.556 | 0.324 | 0.667 | 0.147 | 0.556 | 0.765 | 0.630 | 0.353 | 0.667 |
| Lateral temporal | 86 | 54 | 0.442 | 0.463 | 0.163 | 0.148 | 0.733 | 0.722 | 0.349 | 0.444 | 0.221 | 0.444 | 0.070 | 0.093 | 0.791 | 0.722 | 0.302 | 0.389 |
| Insula | 11 | 6 | 0.364 | 0.333 | 0.182 | 0.000 | 0.727 | 0.833 | 0.273 | 0.500 | 0.364 | 0.333 | 0.000 | 0.000 | 0.545 | 0.500 | 0.182 | 0.167 |
| Paracentral | 29 | 16 | 0.414 | 0.438 | 0.138 | 0.188 | 0.655 | 0.500 | 0.241 | 0.563 | 0.172 | 0.688 | 0.103 | 0.188 | 0.586 | 0.750 | 0.138 | 0.438 |
| Orbitofrontal | 23 | 6 | 0.739 | 1.000 | 0.435 | 0.167 | 0.783 | 1.000 | 0.565 | 0.167 | 0.261 | 0.833 | 0.087 | 0.333 | 0.870 | 0.833 | 0.478 | 0.500 |
| Prefrontal | 49 | 14 | 0.531 | 0.714 | 0.367 | 0.286 | 0.694 | 0.857 | 0.449 | 0.571 | 0.367 | 0.714 | 0.163 | 0.571 | 0.694 | 0.857 | 0.490 | 0.714 |
| Cingulate | 13 | 6 | 0.385 | 0.500 | 0.385 | 0.333 | 0.692 | 0.667 | 0.692 | 0.833 | 0.385 | 1.000 | 0.308 | 0.500 | 0.923 | 0.667 | 0.538 | 0.833 |
| Total# Ch. pairs across NC | 365 | 221 |  |  |  |  |  |  |  |  |  |  |  |  |  |  |  |  |
| Average proportions |  |  | 0.402 | 0.463 | 0.229 | 0.171 | 0.638 | 0.701 | 0.341 | 0.513 | 0.274 | 0.578 | 0.124 | 0.288 | 0.702 | 0.646 | 0.350 | 0.565 |
| Standard deviations |  |  | 0.150 | 0.230 | 0.111 | 0.101 | 0.162 | 0.151 | 0.174 | 0.159 | 0.086 | 0.258 | 0.075 | 0.194 | 0.144 | 0.151 | 0.126 | 0.193 |

**Table 6.**
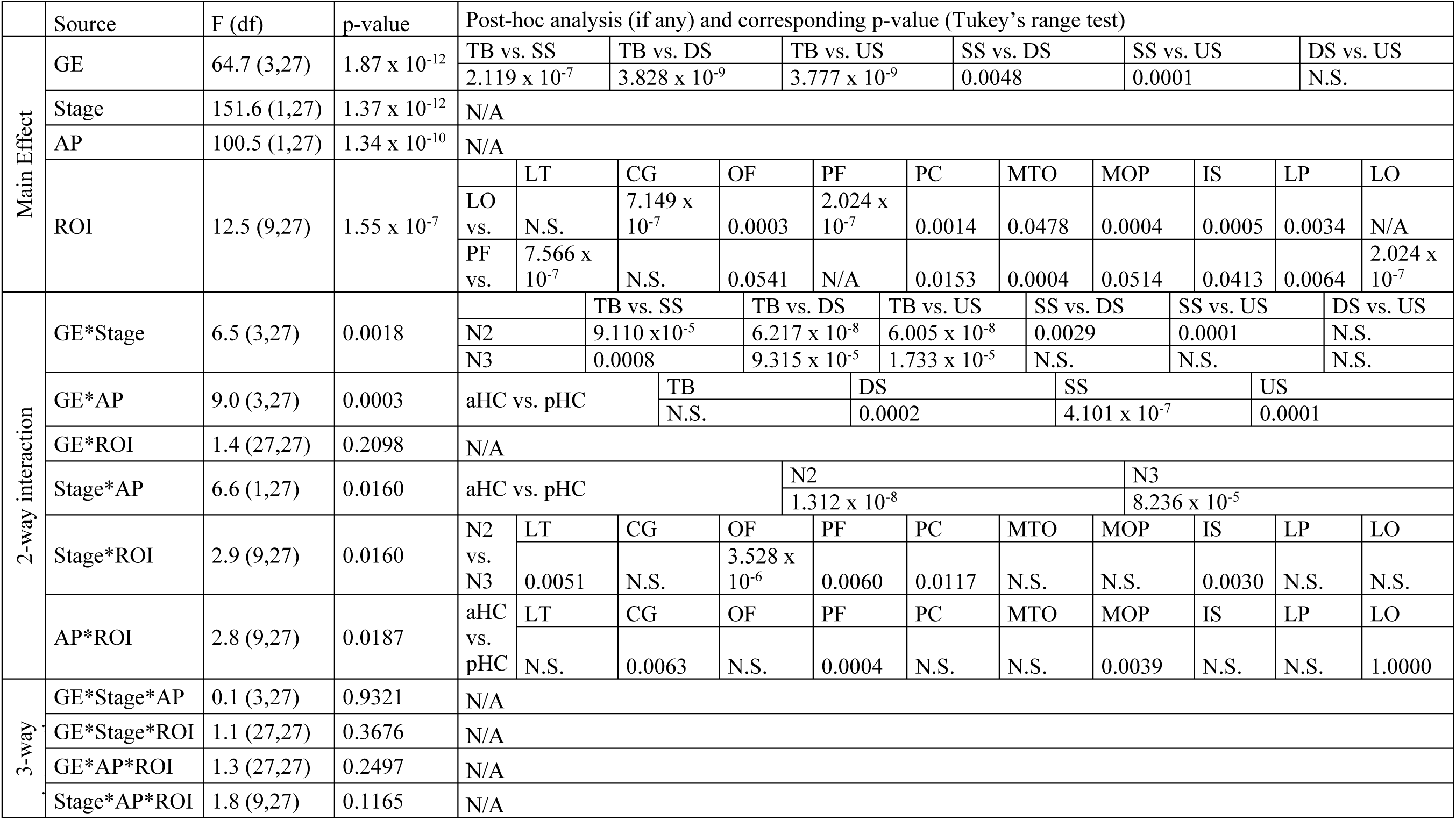
Summary of 4-way ANOVA comparing associations between HC-SSR and NC-GE. For each HC-NC electrode pair, and sleep stage (N2 versus N3), the significance of the NC-GE occurrence times relative to the HC-SSR center was determined (see Fig. 3*A* and Methods). We then tallied the proportion of significant channel pairs according to their HC location (aHC, pHC), NC location (see Table 4-1 and Fig. 5*A*), and GE type (TB, DS, SS, US). These proportions were then entered into ANOVA. LO: lateral occipital; LT: lateral temporal; CG: cingulate; OF: orbitofrontal; PF: prefrontal; PC: paracentral; MTO: medial temporo-occipital; MOP: medial occipito-parietal; IS: insula; LP: lateral parietal. N.S.: not significant (for post-hoc Tukey’s range tests, p > 0.1). df: degrees of freedom.

**Table 7.**
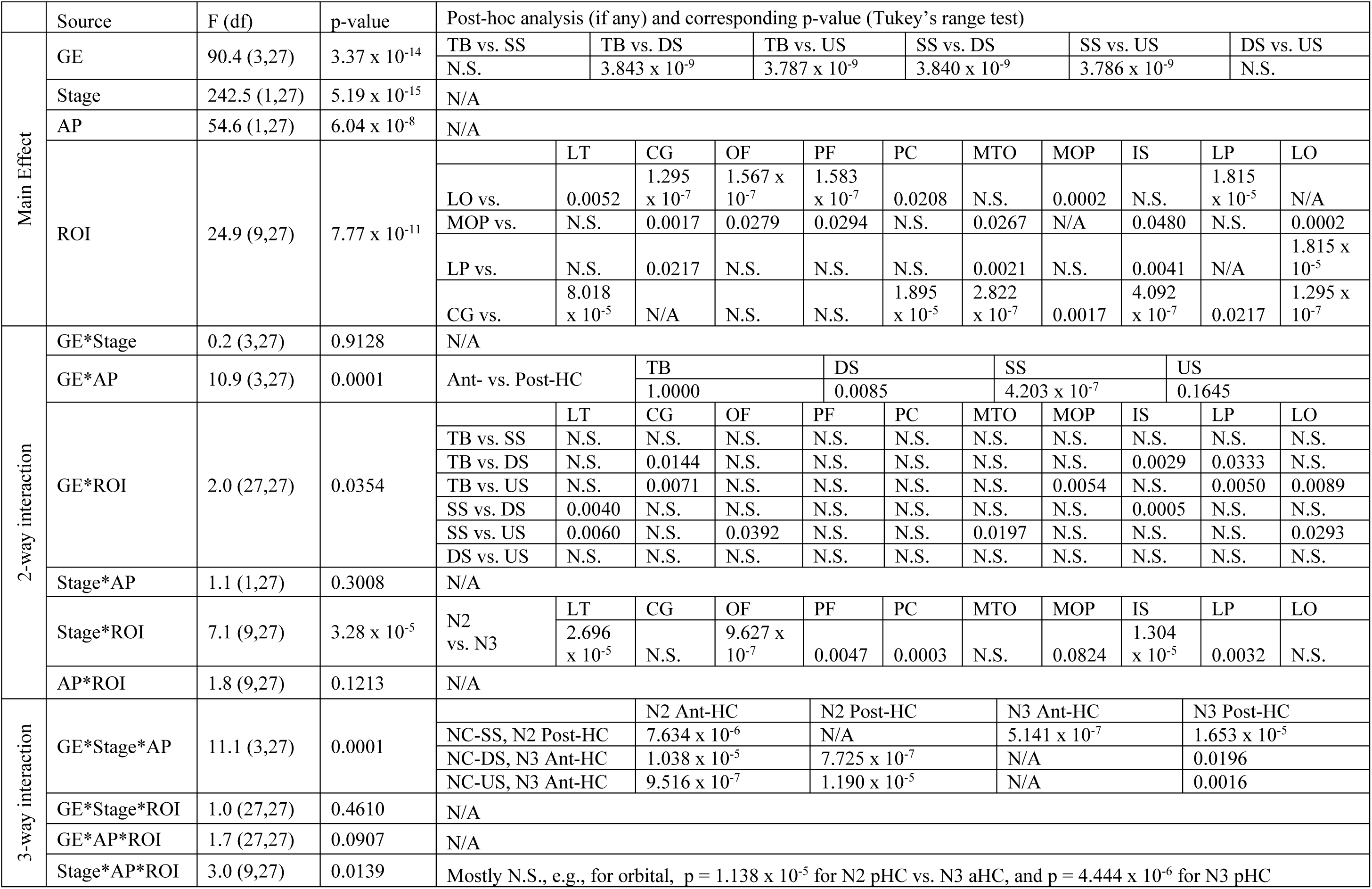
Summary of 4-way ANOVA comparing associations between HC-SS and NC-GE. For each HC-NC electrode pair, and sleep stage (N2 versus N3), the significance of the NC-GE occurrence times relative to the HC-SS onset was determined (see Fig. 3*B* and Methods). We then tallied the proportion of significant channel pairs according to their HC location (aHC, pHC), NC location (see Table 4-1 and Fig. 5*A*), and GE type (TB, DS, SS, US). These proportions were then entered into ANOVA. LO: lateral occipital; LT: lateral temporal; CG: cingulate; OF: orbitofrontal; PF: prefrontal; PC: paracentral; MTO: medial temporo-occipital; MOP: medial occipito-parietal; IS: insula; LP: lateral parietal. N.S.: not significant (for post-hoc Tukey’s range tests, p > 0.1). df: degrees of freedom.

Similarly, given the lateralization of human hippocampal function, we were interested in examining whether ipsilateral HC-NC channel pairs would show different relationships from contralateral pairs across GE types, NREM stages, or NC ROIs. Due to a sparse representation of contralateral channel pairs for individual ROIs, we calculated proportions for larger regions by combining both HC and NC sites: aHC and pHC channels were combined into a single HC category, and the 10 NC ROIs used in previous analysis were collapsed into two: “fronto-central” and “non-frontal” (Table 4-1). We then computed for each ROI the proportion of NC channels significantly coupled to HC-SS/SSR (Table 8), and performed 4-way ANOVA as previously described, with the ipsilateral/contralateral factor replacing the aHC/pHC factor. All post-hoc analyses for both ANOVAs were performed with Tukey’s range test. For each ANOVA, we checked the normality assumption by conducting the Lilliefors test on the residuals.

**Table 8.** Proportions of HC-NC channel pairs with significant GE-SSR or GE-HC SS relationships across NC, separated by ROI, NREM stage, NC-GE type, and ipsilateral/contralateral HC-NC pairings. ipsi: ipsilateral. contra: contralateral.

|  |  | Proportion of channels with significant NC-GE & HC-SS/SSR co-occurrence |  |  |  |  |  |  |  |  |  |  |  |  |  |  |  |
| --- | --- | --- | --- | --- | --- | --- | --- | --- | --- | --- | --- | --- | --- | --- | --- | --- | --- |
|  | NC-GE Type | TB |  |  |  | DS |  |  |  | SS |  |  |  | US |  |  |  |
|  | Stage | N2 |  | N3 |  | N2 |  | N3 |  | N2 |  | N3 |  | N2 |  | N3 |  |
| HC Event | ROI \ SWR Source | ipsi | contra | ipsi | contra | ipsi | contra | ipsi | contra | ipsi | contra | ipsi | contra | ipsi | contra | ipsi | contra |
| SSR | frontocentral | 0.206 | 0.133 | 0.099 | 0.133 | 0.518 | 0.467 | 0.234 | 0.333 | 0.362 | 0.200 | 0.255 | 0.200 | 0.553 | 0.467 | 0.270 | 0.200 |
|  | nonfrontal | 0.169 | 0.138 | 0.052 | 0.049 | 0.436 | 0.228 | 0.225 | 0.179 | 0.300 | 0.260 | 0.169 | 0.098 | 0.495 | 0.382 | 0.228 | 0.171 |
| SS | frontocentral | 0.560 | 0.467 | 0.305 | 0.267 | 0.716 | 0.600 | 0.475 | 0.467 | 0.433 | 0.333 | 0.206 | 0.267 | 0.738 | 0.800 | 0.461 | 0.400 |
|  | nonfrontal | 0.381 | 0.358 | 0.179 | 0.114 | 0.704 | 0.569 | 0.404 | 0.317 | 0.375 | 0.317 | 0.169 | 0.130 | 0.710 | 0.618 | 0.453 | 0.374 |

### Code Accessibility

All custom scripts would be available upon request by contacting the corresponding author and would be delivered through the UCSD RDL-share service.

## Results

In this paper, we quantified HC-SS and HC-SSR in humans, their relationship to HC-SWR and NC-GE, and their variation within different HC regions and sleep stages. We confirmed that ripples phase-modulated by local SS (i.e., HC-SSR) do occur in the human hippocampus, but they are preferentially located in the posterior HC sites. Like HC-SWR (Jiang et al., submitted-1), HC-SS/SSR were strongly related to the NC-GE in a spatiotemporally diverse manner, with particularly strong relationship in stage N2 between posterior HC-SS/SSR and NC-TB, DS, SS and US. The relative latencies are consistent with the order we have established in earlier studies of NC and thalamic GE (Mak-McCully et al., 2017; Gonzalez et al., 2018), i.e. TB→ DS→SS/US, and HC-SS tends to coincide with NC-SS. Preferential co-occurrence, strong coherence, and phase-locking were noted between posterior HC-SS and NC-SS in the posterior parietal and retrosplenial cortices.

### Characterization of human HC-SS and SSR in NREM

We identified human HC-SWR, SSR and SS from intracranial recordings and aligned them with simultaneously recorded cortical events in NREM sleep, in order to determine if their occurrences are related and have a consistent order, and whether their relations change between NREM stages N2 and N3, or between anterior and posterior HC sites. Morphologically normal ripples were isolated from >24 h continuous recordings in 20 stereoelectroencephalography (SEEG) patients with intractable focal epilepsy, with anterior (in 17 patients) and/or posterior HC contacts (in 11 patients), and related to sleep NC-GE in 12-30 bipolar recording channels per patient. Some ripples were found to co-occur with classical sharpwaves (Fig. 1*A1*, with across-patient mean LFP peak-to-peak amplitude of the sharp wave at 127.1 μV), and others did not (Fig. 1*A2, A3*, with peak amplitudes of 8.284 and 14.79 μV, respectively). The latter could sometimes be observed as HC-SSR nestled within HC-SS (Fig. 1*B-C*), as previously reported (Staresina et al., 2015). Time-frequency analysis triggering on HC-SWR ripple centers, SSR ripple centers (with all ripple centers found in the same HC-SS included as separate trials), and HC-SS onsets (Fig. 1*D*) showed that, while HC-SWR tended to be surrounded by a broad-band power decrease, HC-SSR and HC-SS did not. Similar to NC-SS and HC-SWR, HC-SSR and HC-SS in humans were mostly absent in waking (Fig. 1*F*, Fig. 1-1*E*). Notably, HC-SSR and HC-SS did not appear to concentrate in NREM to the same extent as SWR, in agreement with a recent report on human HC-SS in REM (Lestra et al., 2018). However, whether these putative HC-SS hits in Wake/REM are truly HC-SS would be difficult to determine based on our NREM-based detection criteria, and further exploration of this phenomenon would be outside the scope of our current study.

We also found that, while HC sites produce both HC-SWR and HC-SS/HC-SSR (Table 2), the occurrence rates (#/min) of HC-SSR were remarkably lower than those of HC-SWR in aHC (p = 1.602 × 10^-7^ in N2, p = 7.682 × 10^-8^ in N3; two-tailed paired t-tests); no similar difference was observed for pHC, however (p = 0.4911 in N2; p = 0.8038 in N3; two-tailed paired t-tests) (Fig. 1*E*, Fig. 1-1*A-D*). To verify that the relative imbalance of aHC SSR and SWR could not be explained by inter-patient variance alone, we built linear mixed effect models with HC-GE occurrence rates as response variable, and GE type (SWR or SSR), aHC/pHC, and NREM stage (N2/N3) as categorical predictor variables, with patient identity as random effect. We found that, indeed, there is a significant interaction effect of GE type * aHC/pHC on the occurrence rate of HC-GE (p = 7.246 × 10^-7^).

HC-SS/SSR tended to occur more often in pHC than in aHC (p = 0.0018/0.0002 in N2, p = 0.0027/0.0004 in N3, two-tailed two-sample t-tests) (Fig. 1*E*, Fig. 1-1*A-D*). Transitional forms (“SXR”) that were detected as both SWR and SSR constituted a minority of SSR (21.4% in N2 aHC, 26.9 % in N3 aHC, 9.7% in N2 pHC, 13.0% in N3 pHC), and the proportion of SXR among SSR was greater in aHC than in pHC (p = 0.0045 for N2, p = 0.0060 for N3, two-tailed two-sample t-tests), in line with the SWR rate reduction in pHC. Notably, while cortical spindles tend to occur more often in N2, we observed an increase of pHC-SS/SSR in N3 (p = 0.0061 and 0.0088), but not for aHC-SS/SSR (p = 0.0857 and 0.0752; two-tailed paired t-tests). To verify that differences in HC-SS/SSR occurrence rates across hippocampal origin and/or NREM stages could not be explained by inter-patient variance alone, we built linear mixed effect models with HC-GE occurrence rates as response variable, with aHC/pHC and NREM stage as categorical predictor variables and patient identity as random effect. As expected, hippocampal longitudinal origin showed a significant effect on HC-GE rate (p = 0.0003 and 1.925 × 10^-5^ for SSR and HC-SS), while the effect of NREM stage on HC-GE rate was non-significant (p = 0.4785 and 0.2522 for SSR and HC-SS). While there was no significant interaction effect between aHC/pHC and NREM stage for HC-SSR (p = 0.1349), HC-SS did show a significant interaction (p = 0.0324).

NREM stage N3, associated with DS and US (which are in turn associated with SWR), tends to be relatively more concentrated in earlier sleep cycles, whereas stage N2, associated with SS and thus SSR, tends to occur in later cycles. Consequently, we hypothesized that the HC-SSR to HC-SWR ratio would increase over the course of the night. We calculated in each NREM sleep segment the ratio of SSR counts to SWR counts, and compared these ratios in the first and second half of sleep. We found that, while both SWR and SSR occurrence rates decreased in the second half, the extent of the decrease was much more pronounced for HC-SWR (Fig. 1-1*F*). Indeed, the SSR/SWR ratio increased significantly from the first half of NREM sleep to the second half (0.67 vs. 0.88, p = 0.0363, one-tailed paired t-test). However, when we accounted for patient-wise variability by introducing a linear mixed-effect model, we found the effect of early/late NREM on SSR/SWR ratio to be non-significant (p = 0.5667).

Given the presence of both HC-SWR and HC-SSR in the same hippocampus (Table 2), and the preference of HC-SSR to occur in pHC compared to aHC (Fig. 1*E*), we investigated whether a consistent relationship exists between SWR and SSR in aHC and pHC. In 2 out of 8 subjects with both aHC and pHC contacts in the same HC, we found significant elevations in pHC-SSR counts at +100 to +500 ms following aHC-SWR triggers (p < 7.811 × 10^-12^ for all panels in Fig. 2*A1* (N2) and Fig. 2-1*F* (N3), one-sample t-test, with SXR excluded), in contrast to the usual ±200 ms co-occurrence of pHC-SWR with aHC-SWR (Fig. 2*A2*, Fig. 2-1*G*).

Since HC-SS could co-occur with multiple ripples in different spindle cycles (Fig. 1*B*), we tallied the numbers of HC ripples within the duration of HC-SS. Up to 9 ripples in N2 (or 10 in N3) could occur within one HC-SS (Fig. 2*B*, Fig. 2-1*H-J*). Given the potential locality of HC ripples (Patel et al., 2013) and the SS phase modulation of high gamma amplitude (Fig. 2*C*), these ripple counts might be underestimates if each spindle cycle could contain a ripple. More than half of the HC-SS overlapped with at least one ripple from the same HC site. Among the population of HC-SS with at least one co-occurring ripple, more than 60% had two or more ripples within the SS duration.

Time-frequency plots locked to HC spindle peaks yielded ripple power increases that appeared modulated by spindle frequency phase (Fig. 1*D*); this was further supported by intrahippocampal phase-amplitude coupling (Fig. 2*C*). The time-course of this coupling was examined by computing event-related phase-amplitude coupling (ERPAC) between spindle (10-16 Hz) phase and ripple (60-120 Hz) amplitude, using previously published methods (Voytek et al., 2013). All HC-SS in NREM across all patients were included in the computation, separated by longitudinal origin (aHC/pHC), with aligned events being HC-SS starts. As we expected from the phase-amplitude coupling measure (Fig. 2*C*), we found significant event-related modulation of hippocampal ripple amplitude by HC-SS phase (Fig. 2*D*), with the effect being much more prominent in pHC (two-tailed paired t-test over mean ERPAC values from 0 to 1000 ms post-trigger, p < 2.20 × 10^-16^, Cohen’s *d* = 1.74).

### Co-occurrence of HC-SS and HC-SSR in anterior and posterior, or left and right HC

Although initial descriptions of SWR in rodents emphasized their co-occurrence throughout the entire extent of both hippocampi (Buzsaki et al., 1992), more detailed studies showed that isolated SWR are more common (Patel et al., 2013). The dominant and non-dominant hippocampal formations of humans are thought to be specialized for verbal and visuospatial memory, respectively (Glosser et al., 1998), which might predict distinct memory traces and thus low bilateral co-occurrence of HC-SS/SSR. Analogously, the extent to which HC-SSR and/or HC-SS co-occur within the same hippocampus and/or bilaterally ought to be investigated.

We found that, for patients (n = 8) with both aHC and pHC contacts in the same HC, the overlap (i.e. co-occurrence within 50 ms) between aHC- and pHC-SSR (with N2 and N3 combined) was strikingly low, at 1.7±2.1%, though still greater than the mean overlap percentage derived from chance (p = 0.0173, one-tailed paired t-test). Similarly, while the overlap between aHC- and pHC-SS was significantly greater than chance (p = 0.0010, one-tailed paired t-test), the mean percentage remains low (3.7±3.2%). Similarly low overlap percentages were found when using data from N2 or N3 alone (Fig. 2-1*A*).

We also evaluated HC-NC GE overlap in patients (n = 4) with bilateral (left and right) HC contacts, placed in either both anterior or both posterior HC. In these patients, co-occurrence of SSR between HC in different hemispheres (with N2 and N3 combined) was minuscule (0.41±0.47%). This was much less than that usually observed between two contacts within the same HC, and not significantly greater than chance-derived overlap percentages (p = 0.1506, one-tailed paired t-test). Co-occurrence of HC-SS was also rare but more frequent than that of HC-SSR (1.0±0.66%). This small percentage nonetheless was significantly greater than chance (p = 0.0062, one-tailed paired t-test). Similar overlap percentages were found when using data from N2 or N3 alone (Fig. 2-1*B*).

Thus, in humans, a small but significant proportion of HC-SSR and HC-SS co-occurs in both anterior and posterior hippocampal regions, and only a minuscule (but still greater than chance for HC-SS) proportion of HC-SS and HC-SSR co-occur in both left and right hippocampi. Furthermore, by varying the co-occurrence criterion range between 25 ms and 5000 ms, we observed greater-than-chance co-occurrence rates for aHC and pHC-SSR/SS (as well as for left and right HC SSR/SS) initially (Fig. 2-1*A-D*), but the actual co-occurrence rate tapered off exponentially (i.e. showed linear decay in log-log plots) towards chance level beyond 1000 ms. Remarkably, the co-occurrence likelihood of aHC+pHC and bilateral HC-SS/SSR remains in an above-chance plateau until the time window in which co-occurrence is measured expands beyond 500-1000 ms for bilateral HC or for a/pHC. Since the typical significant latency (as estimated in Fig. 3) between NC-GE and HC-SS/SSR is comparable, HC-GE from different HC regions could interact with disparate cortical regions at different times.

### Associations of cortical graphoelements with HC-SSR and HC-SS

In order to examine the relationship between HC-SS/SSR and cortical activity, different cortical graphoelements (NC-GE, including theta bursts (TB), downstates (DS), upstates (US), and spindles (SS)) were identified in simultaneous recordings from all lobes. These recording channels were differential derivations between adjacent contacts spanning the cortical ribbon, and were thus assured of being locally-generated. Detection algorithms have all been extensively validated in our previous studies (Mak-McCully et al., 2015, 2017; Gonzalez et al., 2018). To display the regularities in our data, we constructed histograms of the occurrence times of each NC-GE (TB onset, SS onset, DS peak, US peak) in relation to HC event triggers—HC-SWR ripple center, HC-SSR ripple center (multiple ripples within a single HC-SS counted as separate trials), and HC-SS onset—between each pair of HC (n = 32) and NC (n = 366) channels (n = 20 patients). Separate analyses were carried out for stages N2 vs. N3, for aHC vs. pHC, and for HC-NC pairs in the same vs. opposite hemispheres. Detailed breakdown of histogram contents by GE types and HC-SS/SSR sources (N2 vs. N3, aHC vs. pHC, ipsilateral vs. contralateral) can be found in Extended Data Figure 3-1. For both HC-SS and HC-SSR, across all NC-GE types, in both N2 and N3, the HC-NC associations appeared to be more frequently involving pHC than aHC, as reflected in the elevation of NC-GE counts (Fig. 3-1); this agrees with our earlier observation that HC-SS and HC-SSR predominantly occurred in pHC, and is in contrast with the distribution observed for SWR (Fig. 1*E*).

Of the 598 total HC-NC channel pairs, 555 (93%) of all pairs (429 (94%) of the ipsilateral pairs and 126 (90%) of the contralateral pairs) had at least one histogram showing a significant relationship between the HC-SS/SSR and a NC-GE at p<.0.05 post FDR-correction (randomization test, see section “Experimental design and statistical analysis” in Methods), suggesting widespread NC-GE involvement with HC-SS/SSR. The percentages of significant ipsilateral and contralateral pairs were not significantly different (Fisher’s exact test, p = 0.1888). Of the entire set of 4784 histograms involving HC-SSR and NC-GE, 1247 (26%) showed a significant association; for those involving HC-SS and NC-GE, 2033 (43%) showed a significant association, which is significantly greater than the proportion of histograms coupled to HC-SSR (p = 1.385 × 10^-64^, Fisher’s exact test). The significant histograms (examples are shown in Fig. 3) include all types of NC-GE from channels across all cortical lobes. For HC-SSR, the peak of the histogram represented a mean increase of 157%, or median increase of 86.7% above baseline (measured as the mean height of non-significant bins), with standard deviation of 250%, and positive skew value of 6.40. For HC-SS, the peak of the histogram represented a mean increase of 147%, or median increase of 91.3% above baseline, with standard deviation of 220%, and positive skew value of 9.19.

We summarized these data further by constructing histograms of peak latencies from the histograms described above. Specifically, we determined the latency of peak occurrence rate relative to HC-SSR or HC-SS time for each significant HC-NC-GE histogram, and plotted these as a histogram for each NC-GE type, separately for N2 vs. N3 and aHC vs. pHC (Fig. 4, Fig. 4-1, Fig. 4-2). As noted above, N2 and posterior HC sites had more HC-SSR and HC-SS. Results were also summarized by plotting each cortical channel after morphing to a standardized brain (Fig. 4, Fig. 4-1, Fig. 4-2).In the summary histograms, significant temporal relationships were found between HC-SSR and each of the NC-GE types. As shown in Table 3, we tested for the presence of a significantly non-random distribution of NC-GE peak times with respect to HC-SSR (column 4), and found significant results for both N2 and N3, and both aHC and pHC, for NC-SS and NC-US. NC-TB and HC-SSR were significantly related in N2, but not in N3, possibly due to NC-TB being more abundant and more related to NC-DS in N2 (Gonzalez et al., 2018). NC-DS and HC-SSR were significantly related in N2 for both aHC and pHC, but only for pHC in N3, which may reflect the preference of HC-SSR to occur in pHC. Similarly, in the summary histograms, significant temporal relationships were found between HC-SS and each of the NC-GE types. We also tested for the presence of a significantly non-random distribution of NC-GE peak times with respect to HC-SS (column 8 in Table 3); we found significant results for both N2 and N3, and both aHC and pHC, for NC-TB, DS, SS, and US.

We also examined if the peak latency in the summary histograms between HC-SS/SSR and NC-GE differed significantly from zero, and computed both the mean of the included HC-NC channel pairs (columns 5 and 9 of Table 3) and the center of a distribution fitted over the peak latencies (columns 6 and 10 of Table 3). Based on the fitted distribution centers in N2, NC-TB onsets significantly preceded aHC-SSR by 741 ms, NC-DS peaks significantly preceded by 575 ms, NC-SS onsets preceded by 163 ms, and NC-US peaks followed by 93 ms. Generally similar results could be found for pHC-SSR in N2/N3, based on both mean peak latencies and fitted distribution peak centers (Table 3): at least among the NC-GE that preceded HC-SSR, there appeared to be a reliable sequence of TB→DS→SS for both aHC and pHC in N2. Similarly, for pHC in N3, the TB→DS→SS sequence persisted. Finally, while NC-US were significantly coupled to HC-SSR across N2 and N3 for both aHC and pHC, there were no significant biases to precede or follow HC-SSR, indicating that NC-US tended to co-occur with HC-SSR and were likely to be the last in the sequence of NC-GE types we observed, i.e. TB→DS→SS→US. This sequence appears also in relation to HC-SS (columns 8-11 of Table 3). These observations are consistent with the sequence previously established in earlier studies of NC and thalamic GE (Mak-McCully et al., 2017; Gonzalez et al., 2018).

### HC-NC GE co-occurrence patterns are anatomically selective

Having established that SSR in particular HC sites are significantly associated with GE in NC sites, we asked if co-occurring HC-SSR/NC-GE were randomly distributed across NC locations for a given HC site (Fig. 4-4). Across different GE, for ∼70% of the HC sites, their SSR co-occurred with GE in a non-random distribution of NC sites in N2 (chi-square test of homogeneity). Similar results were obtained for HC-SS in N2, but were less striking in N3 (Figs. 4-4, 4-5).

Given that HC-NC co-occurrences are anatomically non-random (i.e., selective), we further determined if they differed between simultaneously recorded aHC and pHC sites by performing Wilcoxon signed rank tests to evaluate whether for a given GE type, the proportions of GEs from significantly coupled NC sites that overlapped with aHC-SSR/SS differ from the proportions for pHC-SSR/SS. Each test required that both members of a HC site pair have a significant chi-square test result from 1. In total, 8 a/pHC pairs were examined. We found that, for all the qualified a/pHC pairs, the distributions of significantly associated NC channels for aHC and for pHC differed significantly (Fig. 4-6). Thus, different anatomical distributions of NC are significantly coupled with simultaneously recorded aHC pHC.

### Differences in the NC-GE to HC-SSR relationship across NC regions, NREM stages, HC sites, and GEs

While the summaries in Fig. 4 and Table 3 gave us an overview of NC-GE and HC-SS/SSR relations, we noted that there appeared to be regional differences across NC in the strength of temporal correlations. We also considered the possibility that “hot spots” may exist in NC and contribute to the widespread positive skewness in the distributions of GE counts across individual HC-NC GE histograms (Fig. 3-1). We therefore explored the spatial distribution of HC-NC relationship further by tallying the proportion of significant HC-NC channel pairs across different NC regions, with respect to different NC-GE types and different HC-SSR/SS sources (Table 4, Table 5). We then performed two 4-way ANOVAs (one for HC-SSR, another for HC-SS) to compare the main effects of NC-GE type (TB, SS, DS, US), NREM stage (N2 vs. N3), aHC or pHC origin of HC-SS/SSR, and NC ROIs (Fig. 5*A*, coverage listed in Table 4-1), as well as the 2-way and 3-way interaction effects. All post-hoc analyses were performed with Tukey’s range test. We checked the normality assumption by conducting the Lilliefors test on the residuals, and the null hypothesis of normality could not be rejected (p = 0.2058 for HC-SSR, p = 0.0935 for HC-SS). Both the ANOVA and post-hoc analyses’ results (F-statistics and p-values) are summarized in Table 6 (for HC-SSR) and Table 7 (for HC-SS).

**Figure 4.**
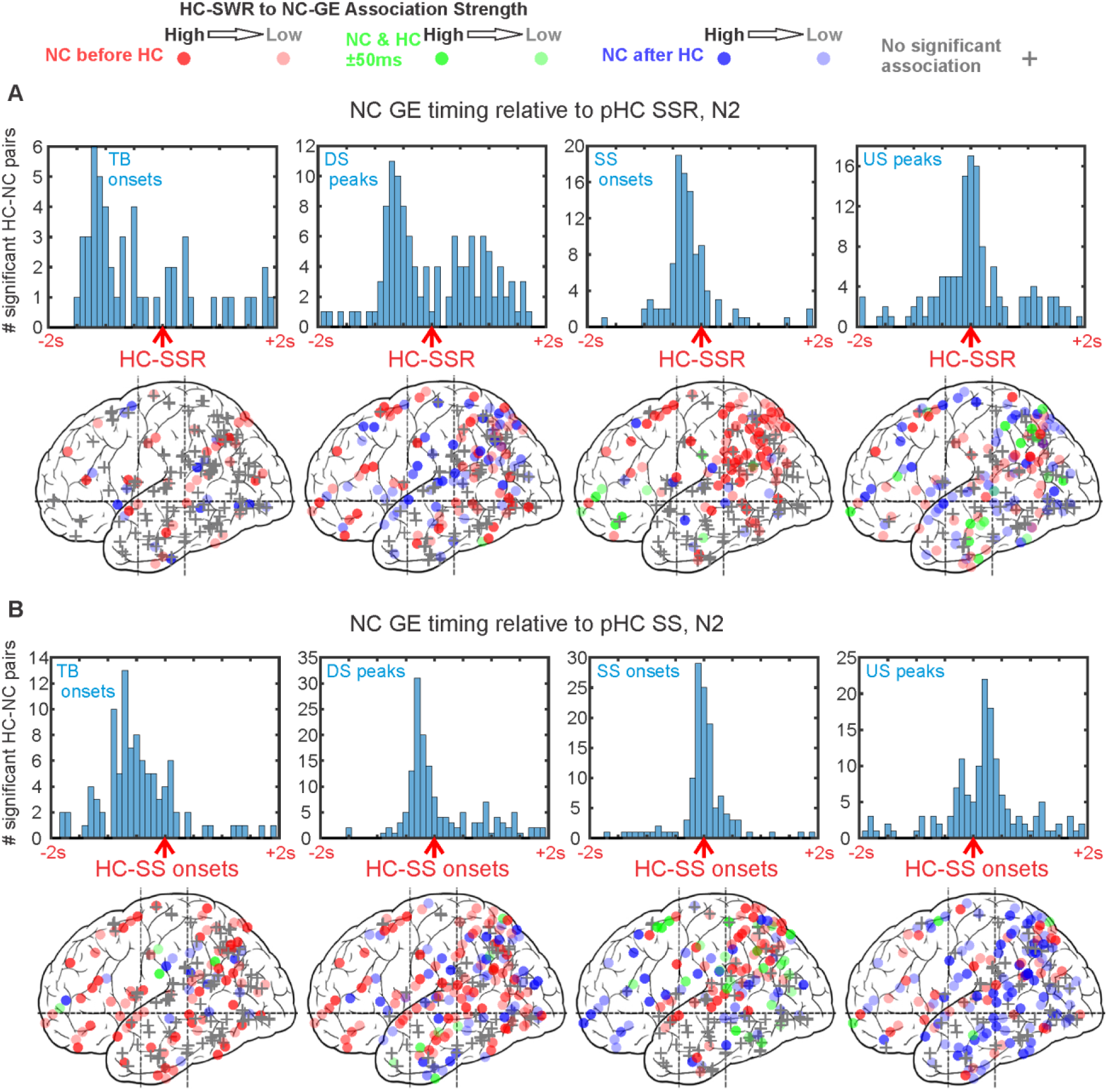
GEs across NC tend to occur with consistent latencies from HC-SSR (***A***) and HC-SS (***B***). ***A***, In N2, with pHC-SSR as reference, NC-GE tend to occur in the following order: TB starts → DS peaks ∼= SS starts → US peaks; NC-US peaks tend to co-occur with HC-SSR. Similar results for pHC in N3 and for aHC in N2/N3 can be found in Extended Data Figure 4-1, and equivalent results for pHC in N2 with only the first SSR in each HC-SS as trigger can be found in Figure 4-3. ***B***. In N2, with pHC-SS as reference, NC-GE tend to occur in the following order: TB starts → DS peaks → SS starts → US peaks. NC-SS tend to co-occur with HC-SS, and widespread NC-US tend to follow. Similar results for pHC in N3 and for aHC in N2/N3 can be found in Figure 4-2. For both ***A*** and ***B***, the top rows contain histograms of peak latencies for HC-NC channel pairs with significant temporal correlations between HC-GE and NC-GE. Each count is the peak latency of a particular HC-NC channel pair. The bottom rows contain maps of peak latency between HC-GE and NC-GE for cortical channels. Circles indicate where NC-GE relationships with HC-SSR/SS are significant, while plus signs represent non-significant channels. The intensity of each circle corresponds to the strength of HC-SSR to NC-GE coupling estimated, and the color of each circle indicates peak latency: red for NC-GE before HC-SSR, blue for NC-GE after HC-SSR, and green for NC-GE co-occurring with HC-SSR (i.e. within 50 ms of each other). Both hemispheres and medial and lateral cortical sites are superimposed in each plot. Additional statistical results on the non-homogeneity of HC-coupled NC site distributions can be found in Extended Data Figures 4-4, 4-5, and 4-6.

**Figure 5.**
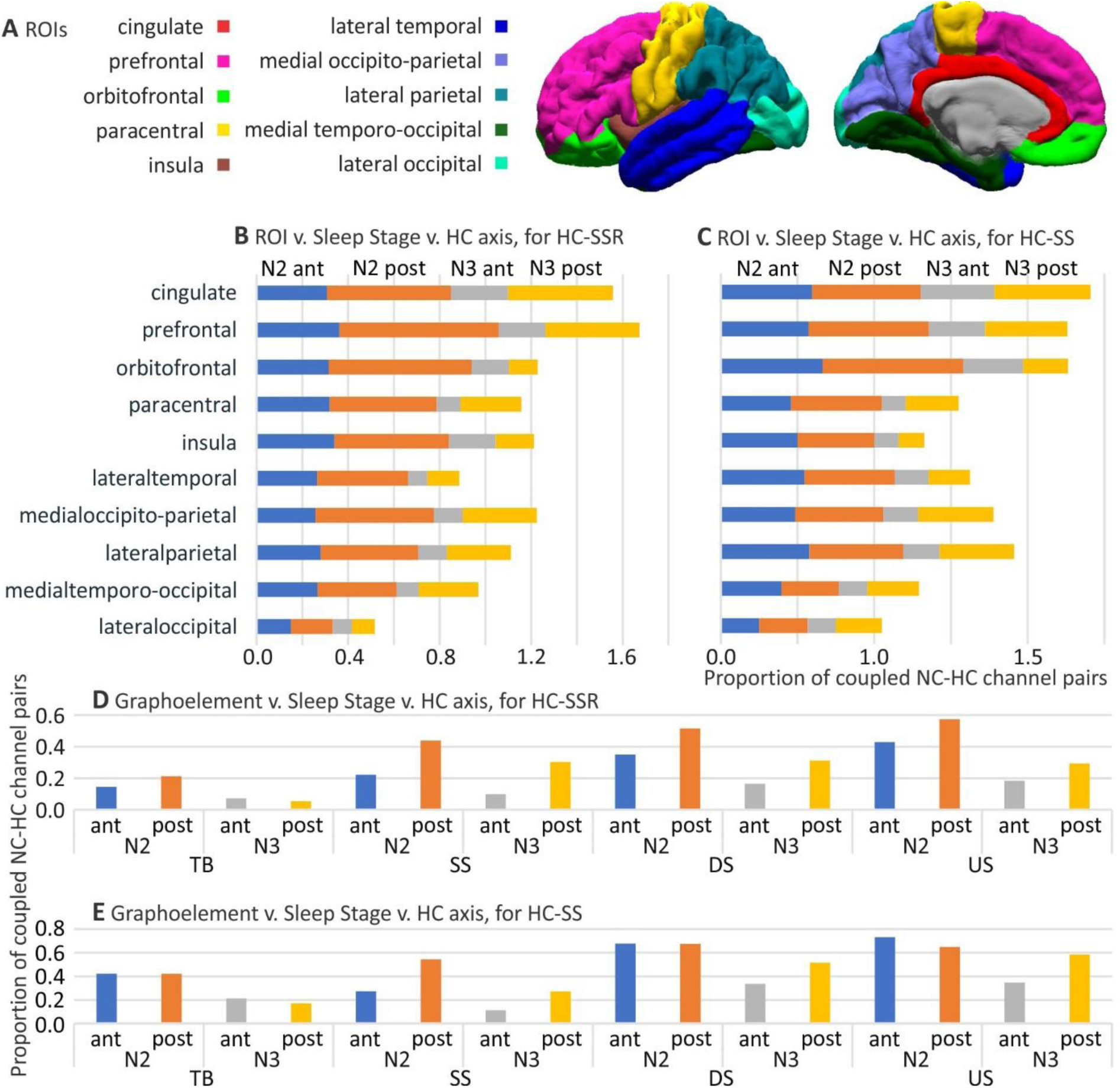
Strength of HC-NC association shows both correlativity and variability across GE types, NREM stages, HC-SS/SSR sources, and NC regions. ***A***. Map of NC ROIs used for analysis (see Table 4-1). ***B***, ***C***. Stacked bar graph of mean significant (evaluated as in Fig. 3) HC-NC channel pair proportions with regard to all channels found in each ROI. Data in ***B*** was obtained by averaging across GE types for each SSR origin (different NREM stages and HC contact positions); similarly, data in ***C*** was obtained by averaging across NC-GE types for each HC-SS origin. ***D***, ***E***. Global (i.e. across all NC ROIs) proportions of significant HC-NC channel pairs for HC-SSR (***D***) and for HC-SS (***E***). Panels ***B***-***E*** share the same color legend. ant: anterior hippocampus; post: posterior hippocampus.

For the ANOVA performed over proportions of NC channels showing significant NC-GE to HC-SSR temporal correlations, all 4 main effects were significant. The main effect for GE was due to the mean proportions of SSR-responsive HC-NC channel pairs differing between all GE type pairs except NC-DS and NC-US. The mean proportions were 0.1252 for TB, 0.2822 for SS, 0.3591 for DS, and 0.3874 for US (Fig. 5*D*). The NREM stage main effect was due to more HC-NC channel pairs being significantly related to HC-SSR in N2 (mean proportion of 0.3784) than in N3 (0.1985). The aHC/pHC main effect was due to more of NC being related to pHC-SSR (0.3617) than to aHC-SSR (0.2152), consistent with our previous observation of higher HC-SSR density in pHC than in aHC. The ROI main effect was due to a stronger HC-NC association by prefrontal locations, followed by cingulate, orbitofrontal, and medial occipito-parietal cortices; in contrast, lateral occipital and lateral temporal regions had the weakest associations, particularly in N3 (Fig. 5*B*, Table 6).

Except for GE * ROI, all 2-way interaction effects from the SSR ANOVA were significant. None of the 3-way interaction effects were significant. The lack of significance for GE * ROI interaction suggests that, overall, the spatial pattern of HC-NC association does not differ across NC-GE types. The GE * NREM stage interaction was due to NC-SS interacting less with HC-SSR than do US and DS, in N2 but not in N3. The GE * aHC/pHC interaction was due to the absence of a pHC advantage for TB in N3. For SS, in particular, pHC proportions were about twice as large as aHC proportions in N2, and about three times as large in N3 (Fig. 5*D*). The NREM stage * aHC/pHC interaction was due to a relatively greater difference in HC-NC coupling by pHC versus aHC during in N2 than N3 (0.1841 vs. 0.1090) (Fig. 5*D*). The NREM stage * NC ROI interaction was due to some locations (orbitofrontal, lateral temporal, prefrontal, paracentral, and insula) showing stronger coupling during N2 than N3, while others (generally more posterior) were about equal (Fig. 5*B*). Finally, the aHC/pHC * NC ROI interaction, was due to a significant aHC/pHC preference limited to cingulate, prefrontal, and medial occipito-parietal regions (Fig. 5*B*), although both aHC- and pHC-SSR appeared to engage the whole cortex (Fig. 3*A*, Fig. 4*A*, Fig. 4-1).

In a companion paper (Jiang et al., submitted-1) we performed a 4-way ANOVA equivalent to the one above, but for HC-SWR. In order to compare the previous results with SWR to the current results with SSR, we included both datasets in a 5-way ANOVA with factors of SWR/SSR, NC-GE type (TB, DS, SS, US), NREM stage (N2/N3), aHC/pHC, and ROI. The normality assumption was checked (p = 0.1478, Lilliefors test on residuals). We found that HC-SWR/NC-GE coupling was more likely than HC-SSR/NC-GE coupling (SWR/SSR main effect, F(1,27) = 117.4, p = 2.471 × 10^-11^). Coupling of SWR/SSR to NC-GE significantly interacted with the type of NC-GE (F(3,27) = 51.57, p = 2.621 × 10^-11^), the NREM stage (F(1,27) = 7.27, p = 0.0119), and whether the SWR/SSR were generated in aHC vs pHC (F(1,27) = 4.57, p = 0.0417). However, the SWR/SSR * NC ROI interaction showed a trend (F(9,27) = 2.11, p = 0.0650). Notably, post-hoc tests revealed that the SWR/SSR * NC-GE interaction effect was due to HC-SSR preferentially coupling with NC-SS (p = 0.0013), while other NC-GE types are more strongly associated with HC-SWR (p = 1.591 × 10^-4^ for TB, p = 6.027 × 10^-8^ for DS, p = 6.616 × 10^-8^ for US). Overall, these results are consistent with the special relationship observed between pHC-SSR and NC-SS.

### Differences in the NC-GE to HC-SS relationship across NC regions, NREM stages, HC sites, and NC-GE types

We repeated the analysis described above for HC-SSR, but for HC-SS, and summarized the results in Table 7. For the ANOVA performed over proportions of NC channels with significant NC-GE to HC-SS temporal correlation, all 4 main effects were significant. The GE main effect was due to a larger proportion of responsive HC-NC channel pairs for DS (0.5484) and US (0.5658) than for TB (0.3165) and SS (0.3160) (Fig. 5*E*). The NREM stage main effect was due to more HC-NC coupling in N2 (0.5507) than N3 (0.3226). The aHC/pHC main effect was due to more HC-NC coupling by pHC (0.4907) than aHC (0.3826), consistent with our previous observation of higher HC-SS density in pHC than in aHC (Fig. 1-1*A-B*, Fig. 1*E*). All the above effects were comparable to their counterparts in the previous ANOVA regarding HC-SSR, with the exception that NC-SS were significantly more related to HC-SSR than NC-TB were, but no such significant difference was observed for HC-SS overall. Thus, HC-SSR could involve a subset of HC-SS with preferential coupling to NC-SS. The ROI main effect was due to greater coupling by NC regions such as the cingulate, versus low levels by other regions (e.g. lateral occipital) (Fig. 5*C*). These results and post-hoc analysis agree with their counterparts from the HC-SSR ANOVA.

The GE * aHC/pHC interaction was due to aHC/pHC differences being confined to DS and SS. For NC-SS, in particular, pHC proportions were about twice as large as aHC proportions in N2, and more than twice as large in N3 (Fig. 5*E*). The GE * ROI interaction was due to a greater coupling of certain NC-GE types in particular ROIs. The insula, for instance, had higher coupling to DS (0.5833) and US (0.3485) than TB (0.2197) and SS (0.1742). The NREM stage * NC ROI interaction a larger coupling in N2 than N3, especially in orbitofrontal and insula ROIs, compared to the small N2/N3 difference in, e.g., cingulate and lateral occipital (Fig. 5*C*). Notably, for both HC-SSR and HC-SS ANOVAs, all ROIs involving occipital cortices (medial temporo-occipital, medial occipito-parietal, and lateral occipital) showed consistent associations with HC across N2 and N3, while orbitofrontal, prefrontal, paracentral, lateral temporal, and insula ROIs showed preferential associations in N2.

The GE * NREM stage * a/pHC interaction is presented in Fig. 5*E*, where e.g. N2 pHC dominated for NC-SS relations. The NREM stage * a/pHC * NC ROI interaction is illustrated in Fig. 5*C*, where N2 pHC proportion was dominant in some ROIs (e.g. orbitofrontal), while others appeared more balanced (e.g. cingulate, lateral occipital). Overall, with minor exceptions, the NC-GE coupling pattern with HC-SS is similar to that of HC-SSR.

### Relationships between NC-GE and HC-SS/SSR in ipsilateral versus contralateral HC-NC channel pairs

Given the well-known lateralization of human hippocampal function (Glosser et al., 1998), we were interested in examining whether ipsilateral HC-NC channel pairs would show different relationships from contralateral pairs across NC-GE types, NREM stages, or NC ROIs. Due to a sparse representation of contralateral channel pairs for individual ROIs, we combined the 10 NC ROIs used in previous analysis into two: “fronto-central”, which included the previous cingulate, orbitofrontal, prefrontal, and paracentral ROIs; and “non-frontal”, which included the previous lateral temporal, medial temporo-occipital, medial occipito-parietal, insula, lateral occipital, and lateral parietal ROIs. We then computed for each ROI the proportion of NC channels significantly coupled to HC-SSR/HC-SS (Table 8) and tallied the numbers of NC channels with or without significant HC-NC associations for each NC-GE type across N2 and N3 (Table 9).

**Table 9.** Numbers and proportions of ipsilateral (ipsi) and contralateral (contra) NC channels showing significant HC-NC GE coupling. Overall, across all GEs, the proportions of significant channels were lower for contralateral than for ipsilateral.

|  | Stage | N2 |  |  |  |  |  | N3 |  |  |  |  |  |
| --- | --- | --- | --- | --- | --- | --- | --- | --- | --- | --- | --- | --- | --- |
|  | (aHC and<br>pHC<br>combined) | ipsi |  |  | contra |  |  | ipsi |  |  | contra |  |  |
| HC<br>Event | NC-GE Type | # sig<br>chan | # nonsig<br>chan | proportion | # sig<br>chan | # nonsig<br>chan | proportion | # sig<br>chan | # nonsig<br>chan | proportion | # sig<br>chan | # nonsig<br>chan | proportion |
| SSR | TB | 81 | 367 | 0.1808 | 19 | 119 | 0.1377 | 30 | 418 | 0.0670 | 8 | 130 | 0.0580 |
|  | DS | 207 | 241 | 0.4621 | 35 | 103 | 0.2536 | 102 | 346 | 0.2277 | 27 | 111 | 0.1957 |
|  | SS | 143 | 305 | 0.3192 | 35 | 103 | 0.2536 | 88 | 360 | 0.1964 | 15 | 123 | 0.1087 |
|  | US | 230 | 218 | 0.5134 | 54 | 84 | 0.3913 | 108 | 340 | 0.2411 | 24 | 114 | 0.1739 |
| Mean over NC-GE types |  | 165 | 283 | 0.3689 | 36 | 102 | 0.2591 | 82 | 366 | 0.1831 | 19 | 120 | 0.1341 |
| SS | TB | 196 | 252 | 0.4375 | 51 | 87 | 0.3696 | 98 | 350 | 0.2188 | 18 | 120 | 0.1304 |
|  | DS | 317 | 131 | 0.7076 | 79 | 59 | 0.5725 | 191 | 257 | 0.4263 | 46 | 92 | 0.3333 |
|  | SS | 176 | 272 | 0.3929 | 44 | 94 | 0.3188 | 81 | 367 | 0.1808 | 20 | 118 | 0.1449 |
|  | US | 322 | 126 | 0.7188 | 88 | 50 | 0.6377 | 204 | 244 | 0.4553 | 52 | 86 | 0.3768 |
| Mean over NC-GE types |  | 253 | 195 | 0.5642 | 66 | 73 | 0.4747 | 144 | 305 | 0.3198 | 34 | 104 | 0.2464 |

We checked the normality assumption for ANOVA by conducting the Lilliefors test on the residuals; while the null hypothesis of normality could not be rejected for HC-SS (p = 0.5000), the null hypothesis was rejected for SSR (p = 0.0030). Nevertheless, we could observe in Table 9 a consistent reduction in the proportions of NC channels showing significant temporal correlation with HC-SSR, by 13-45% for each NC-GE type (separately in N2 or N3) from ipsilateral to contralateral, with NC-TB in N3 showing the least drop and DS in N2 showing the most. Similarly, for HC-SS, we observed a consistent reduction across NC-GE types in both N2 and N3; with the exception of NC-TB in N3 (40% drop for HC-SS vs. 13% for HC-SSR), the ipsilateral-to-contralateral drop (11-20%) in HC-NC association proportions was smaller with respect to HC-SS than to HC-SSR. This suggests that HC-NC communication for spindles preferentially takes place ipsilaterally, and our subsequent phase-amplitude coupling analyses appeared to support this (Fig. 6), as we would describe in detail later in the Results.

**Figure 6.**
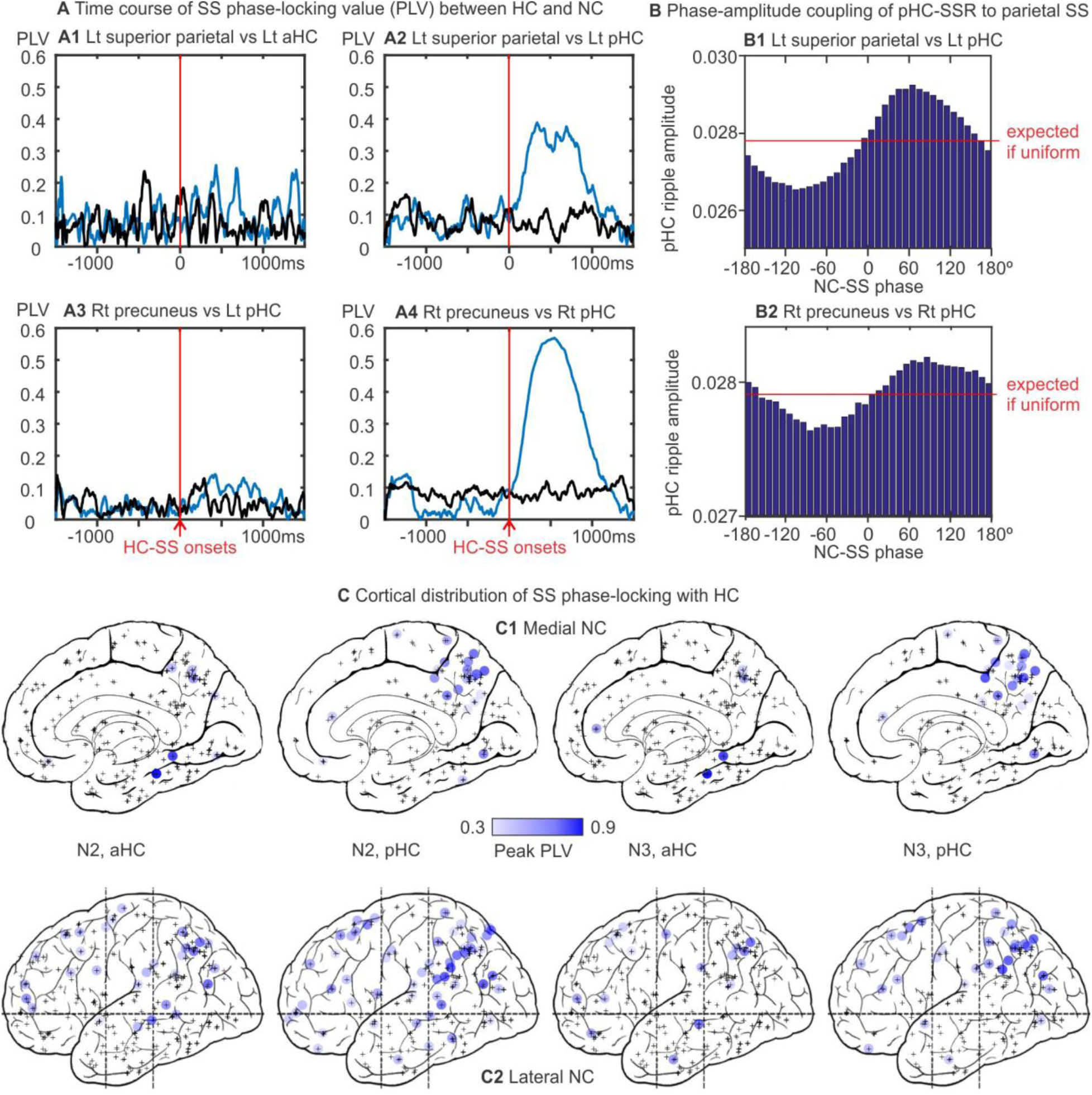
Parietal spindles phase-lock to hippocampal spindles and modulate hippocampal ripple activity. ***A***. HC-SS phase-locking with NC-SS tends to be localized along the longitudinal axis (***A1*** vs ***A2***) and prefers ipsilateral HC-NC channel pairs (***A3*** vs ***A4***). Each panel shows two traces of phase-locking values (PLV); blue trace marks average PLV computed over three-second trials with overlapping HC- and NC-SS, centered on the starts of HC-SS; black trace marks average PLV computed over an equal number of pseudo-trials centered on random time points in NREM (N2+N3). Lt: left; Rt: right. ***B***. Phase-amplitude coupling between the phase of NC-SS (10-16 Hz) in parietal lobe and the amplitude of HC ripple-range activity (60-120 Hz). Each histogram bin covers 10 degrees of phase. The channels in B1 and B2 are the same as those in A2 and A4, respectively. ***C***. 2d projection maps of medial (***C1***) and lateral (***C2***) NC channels showing significant spindle phase-locking with HC. Each blue circle marks a significant NC channel, with color intensity corresponding to peak PLV amplitude. Plus signs mark channels with no significant phase-locking. Projection maps that separate left and right hemispheres can be found in Extended Data Figure 6-1.

We performed 4-way ANOVA as previously described for NC-GE to HC-SS associations, with the ipsilateral/contralateral factor replacing the a/pHC factor. All post-hoc analyses were performed with Tukey’s range test. All 4 main effects were significant, but none of the interaction effects were. As our previous ANOVA already revealed significant main effects for NC-GE type and NREM stage, we focused our attention on the remaining two factors (i.e. ipsilateral/contralateral and combined NC ROI), which provided results beyond those explored in the previous ANOVAs. The main effect of ipsilateral/contralateral channel pair types yielded F = 12.55, p = 0.0383. and together with mean proportions of 0.454 for ipsilateral and 0.400 for contralateral, suggested that more of NC were related to ipsilateral HC-SS than to contralateral HC-SS, although the difference was modest. The main effect of ROIs yielded F = 28.90, p = 0.0126. Since only two ROIs were included in this ANOVA, this result suggested that the more frontal/cingulate parts of NC were significantly more likely to show significant associations between NC-GE and HC-SS than the posterior regions (mean proportion of 0.468 for fronto-central versus 0.386 for non-frontal).

### HC-SS phase-lock to NC-SS, and NC-SS modulate HC ripple activity

A previous report of human HC-SS suggested that scalp-SS could phase-lock with HC-SS, which modulated ripple activity (Staresina et al., 2015). We observed earlier in the results that hippocampal ripples could indeed co-occur with and appeared modulated by HC-SS (Fig. 2*C-D*), and we found widespread NC channels showing significant HC-NC SS temporal correlations, particularly in N2 with respect to pHC (Fig. 4*B*, Fig. 5*E*). To further evaluate spindle-mediated HC-NC coupling, we computed phase-locking values over 3-second trials where NC-SS overlapped with HC-SS for at least one cycle (examples shown in Fig. 6*A*). After PLV computation, for each non-overlapping 50 ms time bin, a two-sample t-test was performed between the actual PLV and the baseline estimate, with the resulting p-values undergoing FDR correction. A given HC-NC channel pair would be considered significantly phase-locking if: 1, more than 40 trials were used in the PLV computation, since small sample size is known to introduce bias (Aydore et al., 2013); 2, at least 3 consecutive time bins yield post-FDR p-values below 0.05. Since HC-SS modulates ripple activity, NC-SS phase-locking with HC-SS would modulate HC ripples by inference; we nonetheless also performed phase-amplitude coupling (PAC) analysis between NC-SS phase and HC ripple amplitude in HC-NC channel pairs with significant SS phase-locking (examples shown in Fig. 6*B*), and estimated the significance of observed PAC by computing average Modulation Indices across trials (each with the same HC-NC SS overlap criterion as used for PLV) and comparing them to bootstrapped baselines (Tort et al., 2010).

We found that significant HC-NC SS phase-locking (over 10-16 Hz, with maximum PLV ranging from 0.3 to 0.9) could be found across widespread NC regions (Fig. 6*C*; Fig. 6-1). Maximum PLV tended to be reached at ∼300-700 ms after HC-SS onsets. We observed a higher density of NC channels showing significant PLV with pHC than with aHC. In particular, 37% of NC channels phase-locked with HC, and out of this channel population, 67% phase-locked with pHC (vs. 33% with aHC) in N2, and 70% phase-locked with pHC (vs. 30% with aHC) in N3. For both N2 and N3, this pHC preference among NC channels with significant SS PLV was significant (Fisher’s exact tests, p = 4.991 × 10^-9^ for N2, p = 2.377 × 10^-7^ for N3). While HC-NC SS phase-locking was observed across NC, only a small proportion had sustained large PLV (e.g. Fig. 6*A2, A4*). Specifically, only 5% of all NC channels had a peak HC-NC SS PLV value above 0.4; in this small population of high PLV channels, ∼70% were parietal (7 in precuneus, 5 in angular gyrus, 4 in intra-parietal sulcus). In addition, for patients with both aHC and pHC contacts in the same hippocampus, NC channels phase-locked to pHC tended not to show significant phase-locking with aHC as well (e.g. Fig. 6*A*), suggesting that HC-NC coupling mediated by spindle phase could be strongly localized along the HC longitudinal axis, which appears in agreement with the rare overlap between aHC- and pHC-SSR/SS reported in the beginning of the Results section.

Similar to the result above with high PLV channel pairs being a small subset of all channels with significant PLV, NC-SS phase to HC ripple amplitude PAC was only observed in about 1/4 of all channels with HC-NC-SS phase-locking. Again, these channels were mainly (∼70%) parietal (6 in precuneus, 3 in angular gyrus, 3 in intra-parietal sulcus). Interestingly, only half of the high SS PLV channel pairs showed significant NC-SS to HC ripple PAC also.

Remarkably, for the population of NC channels from the 8 patients with multiple HC contacts in the same hippocampi, 15 out of the 16 (94%) NC channels showing significant SS phase-locking with aHC also phase-lock with pHC significantly in N2, while conversely only 15 out of the 43 (35%) NC channels showing significant SS phase-locking with pHC also phase-lock with aHC; in N3, this aHC/pHC imbalance in percentages of concurrently coupled NC sites was preserved (7 out of 10 (70%) with aHC vs. 7 out of 24 (29%) with pHC). Thus, if aHC phase-locks SS with an NC site, then pHC does also, but not *vice versa*.

Strong ipsilateral preference for phase-locked HC-NC channel pairs was also observed in the patients with bilateral pHC contacts, with example PLV plots shown in Fig. 6*A*, where the right precuneus preferentially coupled with right pHC. Within the channel pairs showing significant SS PLV involving aHC, in N2, only 5% were contralateral (thus 95% were ipsilateral); in N3, all of the channel pairs with significant PLV involving aHC were ipsilateral. Similarly, for phase-locking channel pairs involving pHC, 89% were ipsilateral in N2, and 90% were ipsilateral in N3. This strong preference for ipsilateral HC-NC SS phase-locking is larger than the preference for ipsilateral co-occurrence of HC-SS and HC-SSR with NC-GE described above.

## Discussion

This study quantifies the characteristics of ripples which co-occur with hippocampal spindles (HC-SSR) in humans. We previously demonstrated that human sharpwave-ripples (HC-SWR) resemble animal HC-SWR in their density, NREM concentration, waveform, and intrahippocampal topography (Jiang et al., submitted-1). HC-SSR and HC-SWR were distinct in important ways: HC-SSR were dominant in pHC, while HC-SWR were in aHC. Both HC-SSR and HC-SWR were coupled with all types of cortical graphoelements (TB, DS, US, and SS) in widespread bilateral locations (Fig. 7). However, HC-SSR were tightly phase-locked with SS in ipsilateral inferior parietal and precuneate sites, whereas HC-SWR showed a preference for prefrontal, central and cingulate cortices. HC-SSR generally occurred during the NC-SS/US, whereas HC-SWR generally occurred ∼400 ms earlier, during the NC-DS. A single HC-SSR is commonly adjacent to multiple ripples on successive peaks of the same HC-SS, separated by ≥50 ms, whereas single ripples on HC-SWR were separated by ∼5s. These differences suggest that HC-SSR and HC-SWR may potentially play complementary roles in memory consolidation, though further support from additional behavioral studies is needed, as our current study lacks behavioral assessments that could clearly demonstrate the involvement of HC-SS/SSR in memory consolidation.

**Figure 7.**
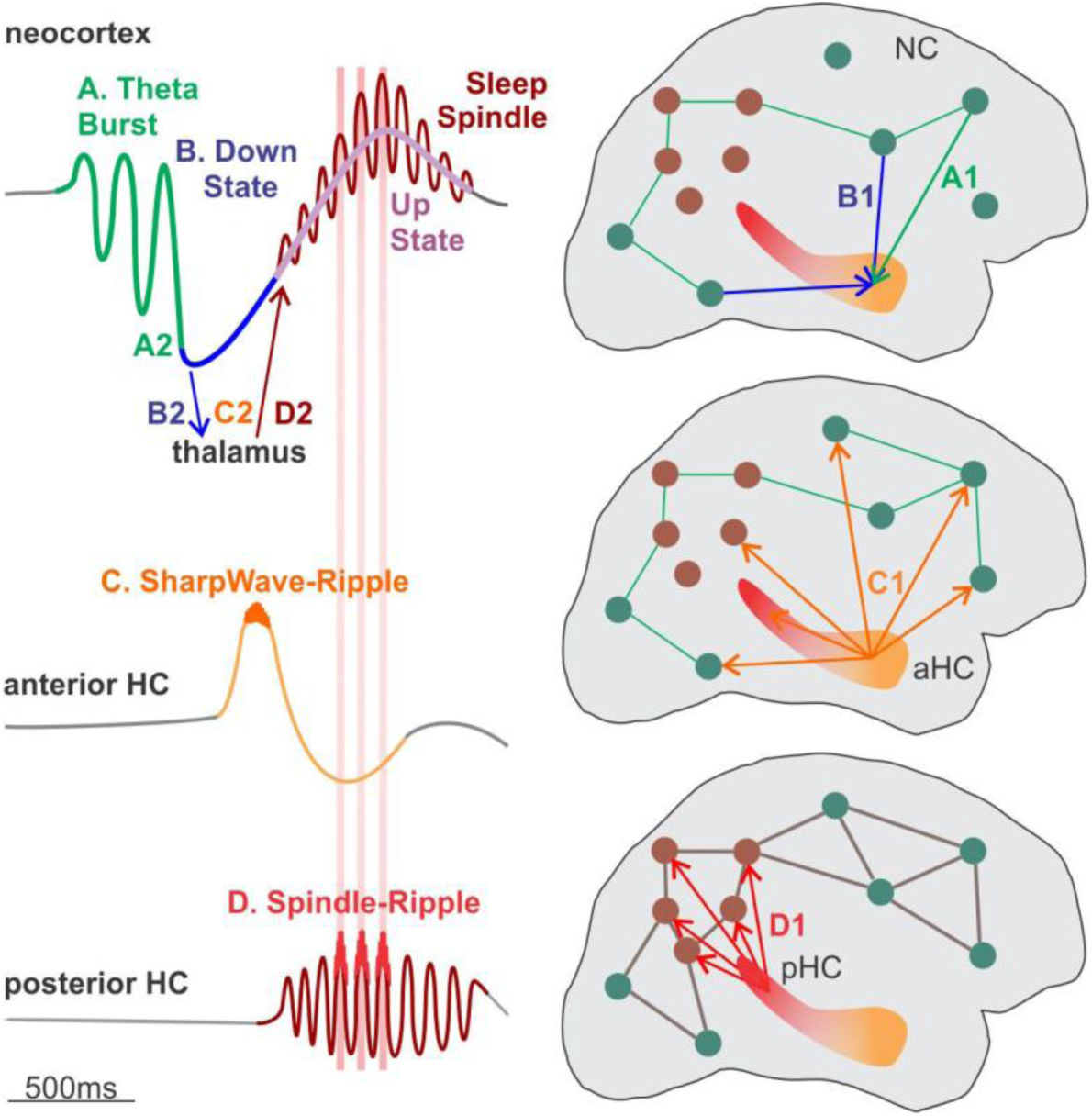
Proposed HC-SWR/SSR interactions with NC-GE in the context of previous findings of NC-HC-thalamic interactions during GE (Mak-McCully et al., 2017; Gonzalez et al., 2018; Jiang et al., submitted-1). ***A.*** The sequence begins with a spontaneous NC-TB. Firing patterns are hypothesized to project to the HC and help inform the context in which they arise (***A1***). Locally, the NC-TB may lead to the NC-DS (***A2***). ***B.*** The NC-DS in turn may contribute to the broadband decrease in HC-LFP preceding SWR (***B1***) and trigger a thalamic-DS (***B2***). ***C.*** The HC-SWR occurs at this critical juncture (Jiang et al., submitted-1), contributing recent memory traces to the NC as it reboots from the NC-DS (***C1***); meanwhile, the thalamic-DS enables the h and T currents to generate a thalamic-SS (***C2***). ***D.*** The pHC-SSR follows, perhaps projecting more detailed recent memory information during multiple ripples (***D1***). The pHC-SSR are tightly phase-locked to SS in inferior and medial parietal cortices (this study), projected from the thalamus (***D2***). Increased firing and Ca^++^ influx during the NC-SS and underlying NC-US may provide an ideal cellular environment for memory consolidation. Note that experimental evidence is lacking for pHC-SSR serving as markers for replay, and the evidence for aHC-SWR is mainly from rodents.

Human HC-SS have previously been recorded in NREM (Montplaisir et al., 1981; Malow et al., 1999; Andrillon et al., 2011; Carpentier et al., 2017), and REM sleep (Lestra et al., 2018). We observed SS in HC with no ictal or interictal involvement, and HC-SS are observed in animals (Miyawaki and Diba, 2016), suggesting that they represent healthy activity. Our confirmation of phase-modulation of HC-ripples by HC-SS, and our demonstration of their coordination with NC-GE, provides a possible framework for HC-NC information transfer, as previously suggested (Siapas and Wilson, 1998; Sirota et al., 2003; Staresina et al., 2015) Antony et al., 2018). However, despite our efforts to eliminate contamination by epileptiform events, our study is limited by the possibility of pathological restructuring, and the possible role of SSR in memory replay is unconfirmed in mechanistic studies.

Animal studies have described varying temporal relationships between HC-SWR and NC-SS. The possibility that NC-SS play an initiating role is implied by the observation that SS can reach the hippocampus via entorhinal cortex and phase-modulate HC firing (Sirota et al., 2003; Isomura et al., 2006). However, others found HC-SWR to precede NC-SS (Siapas and Wilson, 1998; Peyrache et al., 2011). This apparent discrepancy could reflect the fact that while HC-SWR tend to precede NC-SS (Jiang et al., submitted-1), HC-SSR tend to co-occur with NC-SS (Fig. 4*A*, Fig. 4-1). Since some of the reports cited above detected high-frequency power without verifying a co-occurring sharpwave waveform, it is possible that they studied a mixed population of SWR and SSR.

Clemens *et al*. (2011) reported that ripples (including both interictal spikes and physiological activity) recorded medially to the parahippocampal gyrus are phase-modulated by parietal scalp spindles. Similarly, Staresina *et al*. (2015) found a 5% increase in HC ripple band power +/-250 ms to scalp spindles. However, the current report appears to provide the first evidence that human intracranially-recorded NC-SS phase-lock with HC-SS and show phase-amplitude coupling with ripples. HC and parietal spindles may become phase-locked via projections from CA1 to area 7a (Clower et al., 2001), thalamo-hippocampal (Amaral and Cowan, 1980), and/or thalamo-cortical projections (Mak-McCully et al., 2017) (Fig. 7*B-D*).

An important consequence of this co-occurrence and phase-locking of HC-SWR with NC-SS is that if HC-SWR mark replay firing, then this information would be projected to NC at a propitious time (Fig. 7*D*). In humans, NC-SS (Hagler et al., 2018), and especially NC-SS/US increase firing (Csercsa et al., 2010; Mak-McCully et al., 2017; Gonzalez et al., 2018). In rodents, NC-SS (Seibt et al., 2017), and especially NC-SS/US (Niethard et al., 2018), are associated with strong Ca^2+^ influx in NC pyramidal cell apical dendrites, and increased plasticity (Steriade and Timofeev, 2003). Behavioral consolidation in rodents depends on NC-SS/US coupling with HC-SWR (Maingret et al., 2016; Latchoumane et al., 2017). Similarly, scalp SS and SS-US complexes are associated with consolidation in humans (Marshall et al., 2006; Mednick et al., 2013; Niknazar et al., 2015).

The distinctive behavioral contributions of ventral/dorsal HC in rodents, or aHC/pHC in humans, have been characterized in multiple ways. In rodents, ventral HC lesions impair contextual emotional learning, corresponding to its amygdala, orbitofrontal, and hypothalamic connections, whereas dorsal lesions impair spatial learning, corresponding to its retrosplenial and parietal connections (Fanselow and Dong, 2010). Although ventral HC neurons also exhibit place fields, they are less frequent and larger than dorsal HC (Strange et al., 2014), and the smaller ventral HC dentate gyrus may support coarser pattern separation (Poppenk et al., 2013). The anatomical contrasts between ventral and dorsal HC in rodents have been confirmed with resting state BOLD connectivity measures in humans (Ranganath and Ritchey, 2012). Functional imaging studies suggest greater aHC involvement in learning the gist of events, and pHC the details, perhaps consistent with the broader encoding by its cells in rodents (Poppenk et al., 2013; Strange et al., 2014).

The preponderance of SSR in dorsal HC has not been reported in rodents. Indeed, SWR are *more* dense and stronger in dorsal than ventral rodent HC (Patel et al., 2013). One possible interpretation is that pHC-SSR may support a faculty that is particularly well-developed in humans. The cortical areas (angular g., precuneus, and retrosplenial cortex) strongly phase-locked with pHC-SSR, as well as the HC itself, are all associated with episodic memory, which Tulving (Tulving and Markowitsch, 1998) has argued is unique to humans. Lesions restricted to the HC but sparing adjoining temporal cortical areas severely impair episodic memory with relatively preserved non-episodic declarative memories (Vargha-Khadem et al., 1997). Similarly, angular gyrus lesions impair recollective experience without necessarily impairing familiarity judgements (Papagno, 2018). Recollective experience is correlated with hemodynamic activation in these areas (Gilmore et al., 2015), as well as unit-increases (Rutishauser et al., 2017), possibly indicating a role in storing details of episodic memories (Rugg and King, 2018, but see Davis et al., 2018). Recollection is also associated with a parietal positive event-related potential, which is dependent on HC integrity (Smith and Halgren, 1989), and correlated across trials with pHC and retrosplenial BOLD activation (Hoppstädter et al., 2015). HC stimulation can induce intense feelings that current experience is being re-lived (*déjà vu*) as well as vivid recollections of previous experiences (Halgren et al., 1978; Curot et al., 2017). These considerations raise the possibility that pHC-SSR are more prominent in humans because they help support autobiographical memory; further investigations with behavioral assessments are necessary to test this hypothesis.

The ratio of hippocampal to neocortical pyramidal cells is ∼1:33 in rats, and ∼1:2246 in humans (Korbo et al., 1990; West and Gundersen, 1990; West et al., 1991; Pakkenberg and Gundersen, 1997). Thus, the HC→NC fanout is extremely broad in humans. Yet, a central feature of HC-dependent memory is its ability to unify arbitrary elements, and this is presumed to require the HC to interact with all or most of the NC (McNaughton, 2010). Indeed, we have shown that the HC-SWR co-occur with NC-SS/US across widespread cortical areas in all lobes (Jiang et al., submitted-1). Furthermore, HC-SWR only occur ∼12 times per minute, and thus most cortical activity would develop in the absence of HC input from any particular location. Nonetheless, HC-SWR tend to occur at the time of the NC-DS, i.e., at the beginning of the NC-SS/US (Fig. 7*B*), and thus can set the stage for subsequent elaboration of the NC↔NC network. Therefore, the wide HC→NC fanout and low HC-SWR density as well as its timing, might be expected to limit the influence of HC input to setting the general context of NC network evolution rather than its precise content (Fig. 7*C*).

In contrast to aHC-SWR, pHC-SSR produce multiple ripple bursts and thus replay patterns to a focal cortical target during the process of NC cell-assembly. Multiple ripples commonly occur over the course of a single pHC-SSR, with their centers separated by ≥50 ms, compared to HC-SWR which are separated by ∼5s. Furthermore, in contrast to the NC-GE co-occurrence of HC-SWR with widespread NC areas, pHC-SSR restrict their very high level of phase-locking during putative replay to a focal target in area 7a. Finally, the pHC-SSR input to NC is sustained during the SS-US when cell-firing dendritic calcium levels are optimal for synapse formation, as opposed to a single punctate input from HC-SWR at the beginning of the SS-US. Thus, while the HC-SWR input appears well-suited to provide guiding input to widespread NC areas comprising the “gist” of a recent event, the pHC-SSR may be better-suited to impress specific details through repeated inputs to focal NC areas (Fig. 7*C-D*).

In summary, during NREM sleep the human hippocampus generates ripples, and may potentially provide specific information to the neocortex, in two distinct LFP and anatomical contexts: aHC-SWR and pHC-SSR. Both are coordinated with thalamocortical waves which occur in a typical order TB→DS→SS/US, but aHC-SWR occur with the DS and pHC-SSR with SS/US (Fig. 7). aHC is anatomically connected with limbic and prefrontal sites, and aHC-SWR are preferentially coupled with prefrontal and cingulate NC-GE (Jiang et al., submitted-1). pHC is connected to parahippocampal, retrosplenial, precuneus, and inferior parietal cortices, and pHC-SSR are strongly phase-locked with these areas. The low rate of aHC-SWR, their broad bilateral anatomical fanout, and the preference of this fanout for fronto-limbic sites are consistent with the contextual contribution sometimes assigned to aHC (or ventral hippocampus in rodents). In contrast, the rapid repeated ripples during pHC-SSR, phase-locked with focal ipsilateral parietal loci, is consistent with a more detail-oriented consolidation process, if indeed, pHC-SSR are found to participate in replay.

## Acknowledgements

This work was supported by the U.S. Office of Naval Research’s Multidisciplinary University Research Initiatives Program (N00014-16-1-2829), the National Institute of Mental Health (RF1 MH117155 and R01 MH111437), and the National Institute of Biomedical Imaging and Bioengineering (R01 EB009282). The authors would like to thank the following for their support: John Gale, Qianqian Deng, Charles Dickey, Darlene Evardone, Zach Fitzgerald, Chris Gonzalez, Don Hagler, Milan Halgren, Erik Kaestner, Rachel Mak-McCully, Adam Niese, Burke Rosen, and T. G. Venti.

## Supplemental Material

**Figure 1-1.**
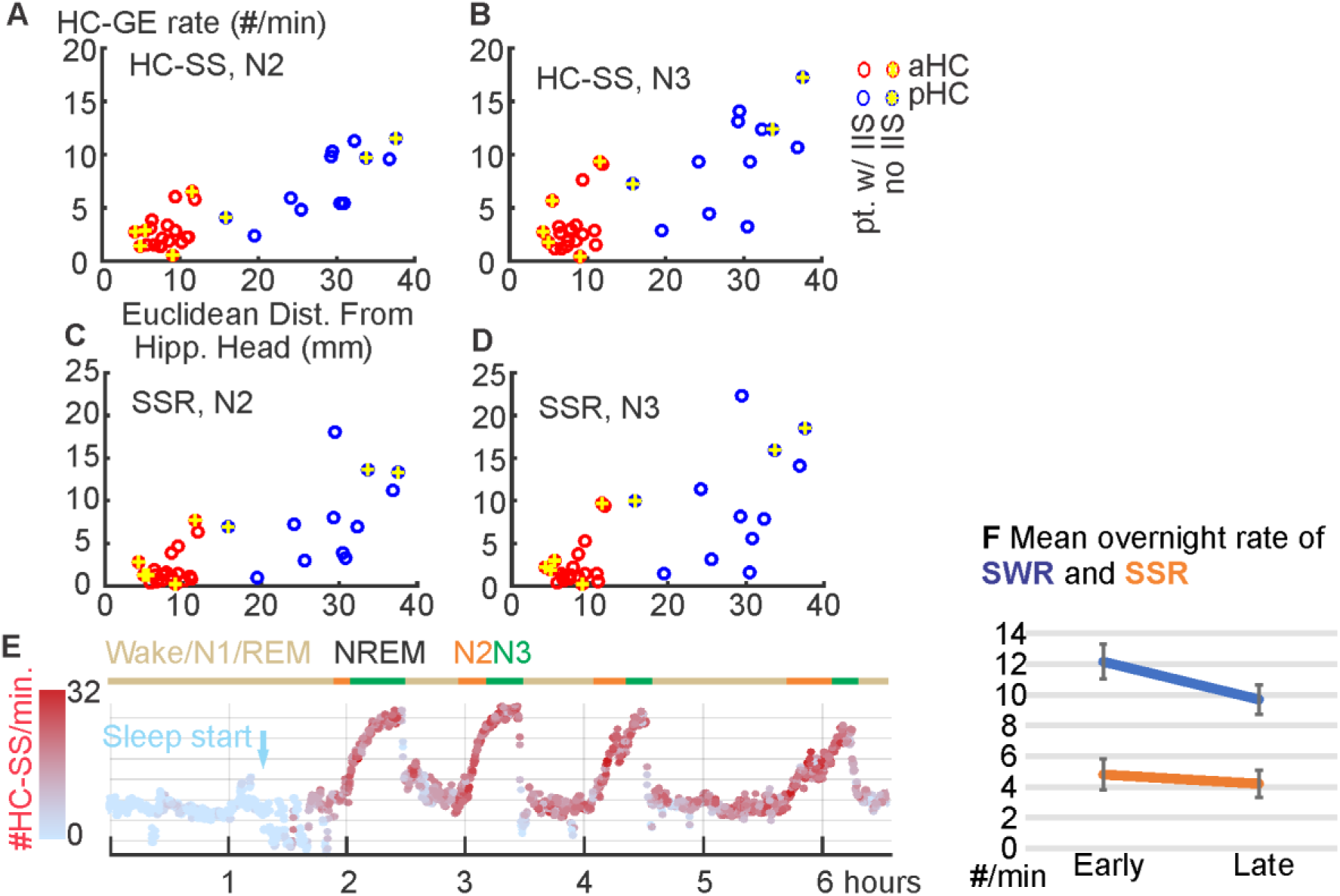
additional information on HC-SS/SSR distributions over the longitudinal axis of HC and over behavioral states. ***A***-***D***. Distribution of HC-SS and HC-SSR occurrence rates in different NREM stages across the HC longitudinal axis. Each circle represents data from one hippocampal contact. Red circles indicate contacts in aHC; blue circles indicate those in pHC. Yellow highlights mark the contacts from interictal-free patients. Pt.: patient. ***E***. Example state plot showing the separation of NREM sleep from waking/REM in ∼8-hour LFP recording and HC-SS occurrence rate (color coded with red intensity) over time. N2/N3 periods are marked with orange/green horizontal lines, respectively. Light blue arrow marks the beginning of NREM sleep. ***F***. Mean rates of HC-SWR and HC-SSR in the first half (“Early”) versus second half (“Late”) of NREM sleep across all nights and patients.

**Figure 2-1.**
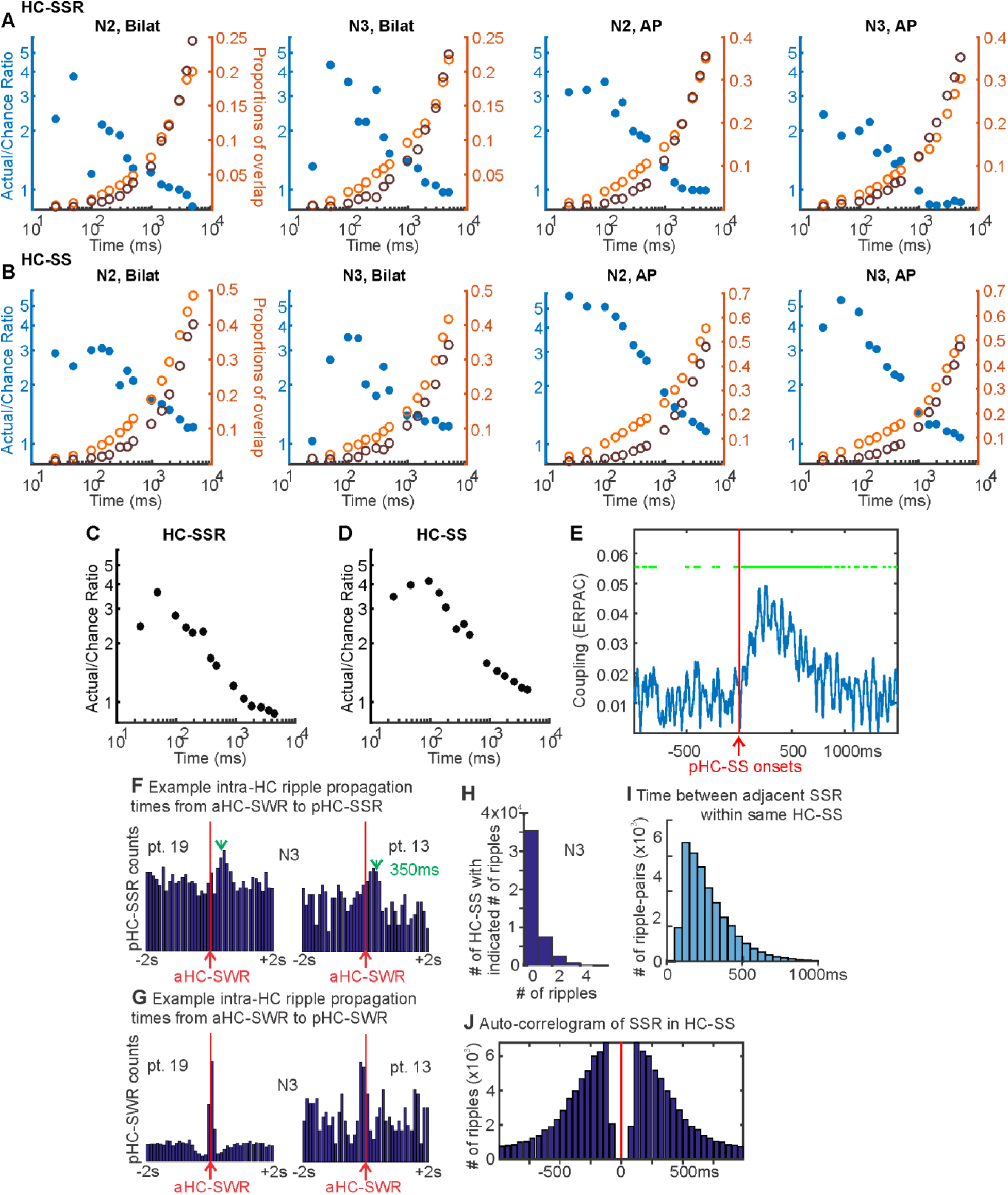
Co-occurrence likelihood distributions over different time windows for bilateral hippocampi and along the HC longitudinal axis. ***A***-***B***. For both bilateral HC site pairs and Ant-HC/Post-HC site pairs, HC-SSR (***A***) and HC-SS (***B***) co-occurrence likelihood approaches chance as time window for co-occurrence expands. For bilateral HC, high variability in the ratio between actual and chance proportions (of overlapping HC-SS/SSR) was observed, likely due to the relatively small sample size. Blue filled circles mark (wrt. left y-axis) the ratios of actual over chance overlap proportions, and orange/brown unfilled circles mark (wrt. right y-axis) the corresponding actual (orange) and chance (brown) proportions. All axes for each panel are logarithmic. ***C***. Combined data from ***A*** show that HC-SSR co-occur between different HC locations (i.e. across both hemispheres or between aHC and pHC of the same hemisphere), at a low rate (<4%) that falls to chance before 1 s of separation. ***D***. Similar findings for HC-SS after combining data from ***B***. ***E***. The time-course of intra-hippocampal event-related phase-amplitude coupling (ERPAC) between spindle (10-16 Hz) phase and ripple (60-120 Hz) amplitude, computed over HC-SS in NREM across all patients for pHC, where pHC events are randomly sub-selected to have the same number of events involved as aHC (Fig. 2*D1*). Events are aligned at HC-SS starts (red vertical lines). Green line segments mark the time segments with significant coupling (p < 0.001). ***F***-***G***, Peri-stimulus time histograms in N3 with aHC-SWR as triggers and either pHC-SSR (***F***) or pHC-SWR (***G***) as tallied events, with HC-SXR excluded from ***F***; pHC-SSR tend to follow aHC-SWR by ∼350 ms (marked with green arrows in A), and aHC/pHC-SWR tend to co-occur within ±200 ms. ***H***. Histograms of ripple counts across all HC-SS in N3. Not shown are the HC-SS with ≥5 ripples (n=211, max 10). In both N2 (Fig. 2B) and N3, only ∼10% of the HC-SS had 3 ripples or more nestled within, possibly owing to less prominent ripples in LFP signal being under the detection threshold. ***I***. Histogram of temporal intervals between adjacent ripples within the same HC-SS, peaking between 100 and 150 ms, with the median of the distribution at 234 ms. Each histogram bin covers 50 ms. The distance between adjacent SSR within the same HC-SS peaks at between 100 and 150 ms, with the median inter-ripple distance being 234 ms. ***J***. Auto-correlogram of ripple occurrence times, triggering on ripples within HC-SS, over ±1000 ms, with 50 ms bins. Red vertical line marks the centers of each SSR. The drop in correlogram counts between 0 and 100 ms agrees with ripple modulation at spindle frequency (Fig. 1C, Fig. 2C-*D*), although the absence of counts between 0 and ±50 ms is partially accounted for due to the ripple detection criteria we used (see “Hippocampal graphoelement selection” in Methods).

**Figure 3-1.**
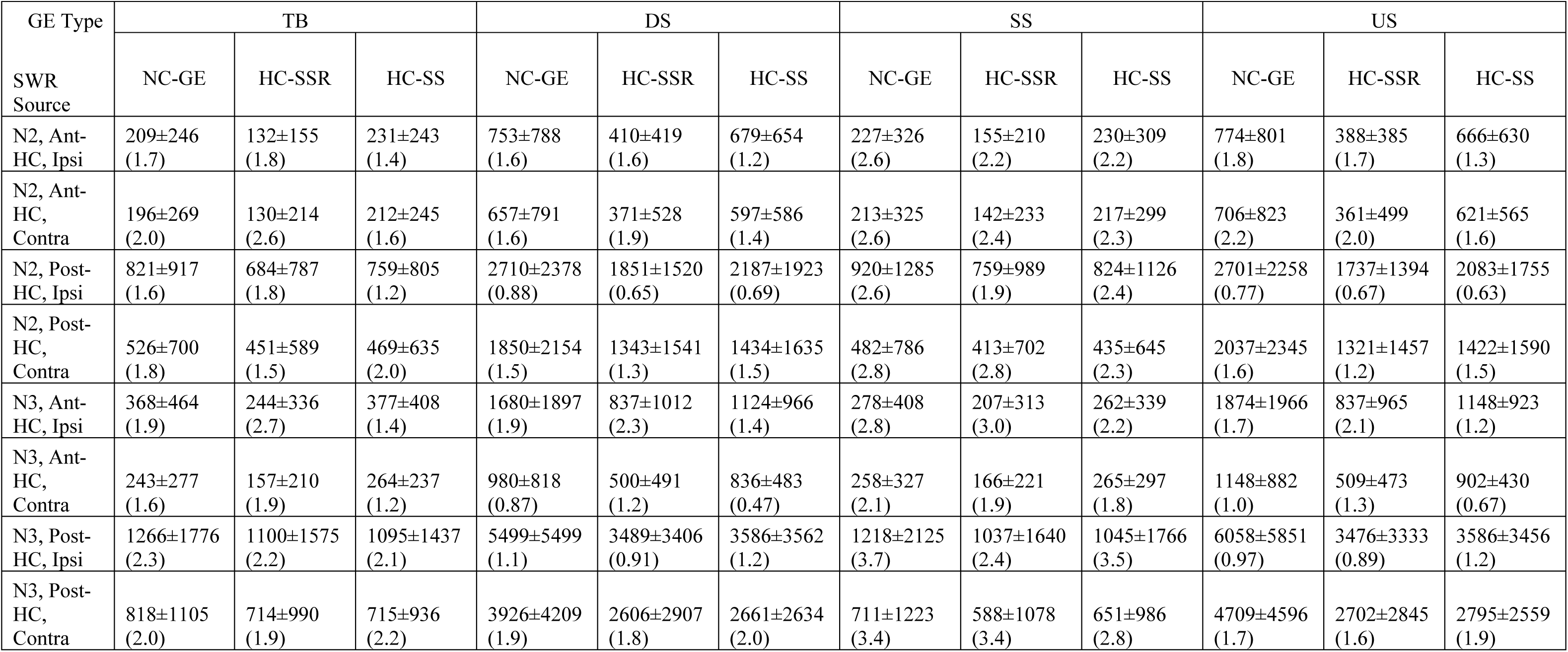
Summary of mean number of events in NC-GE to HC-SSR/SS histograms for individual channel pairs. Each cell is formatted as follows: Mean±Standard Deviation (Skewness). Ipsi: ipsilateral. Contra: contralateral.

**Figure 4-1.**
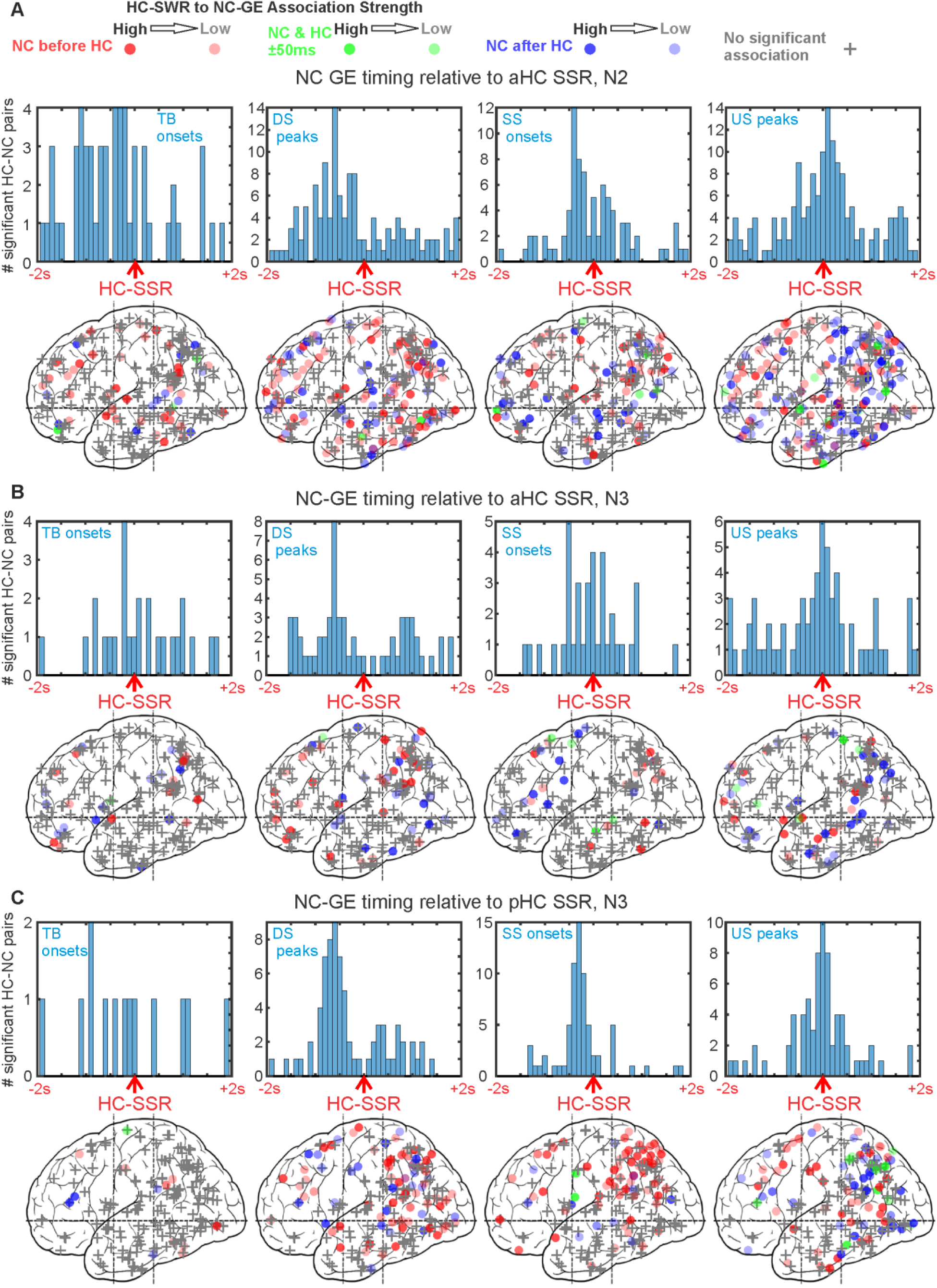
DSs, SSs, and USs across NC tend to occur with consistent latencies from HC-SSR in N3. ***A***-***C***. Top rows: Histograms of peak latencies for HC-NC channel pairs with significant temporal correlations between HC-SSR and NC-GE. Each count is the peak latency of a particular HC-NC channel pair. Bottom rows: maps of peak latency between HC-SSR and NC-GE for cortical channels. Circles indicate where GE-SSR relationships are significant, while plus signs represent non-significant channels. The intensity of each circle corresponds to the strength of GE-SSR coupling estimated (as in Fig. 3), and the color of each circle indicates peak latency: red for NC-GE before HC-SSR, blue for NC-GE after HC-SSR, and green for NC-GE co-occurring with HC-SSR (i.e. within 50 ms of each other). Both hemispheres and medial and lateral cortical sites are superimposed in each plot. ***A***. In N2, with aHC-SSR as reference, NC-GE tend to occur in the following order: TB starts → DS peaks ∼= SS starts → US peaks. ***B***. In N3, with aHC SSR as reference, NC-GE tend to occur in the following order: DS peaks → SS starts → US peaks. ***C***. In N3, with pHC SSR as reference, NC-GE tend to occur in the following order: DS peaks → SS starts → US peaks. In both ***A*** and ***B***, US peaks tend to co-occur with HC-SSR.

**Figure 4-2.**
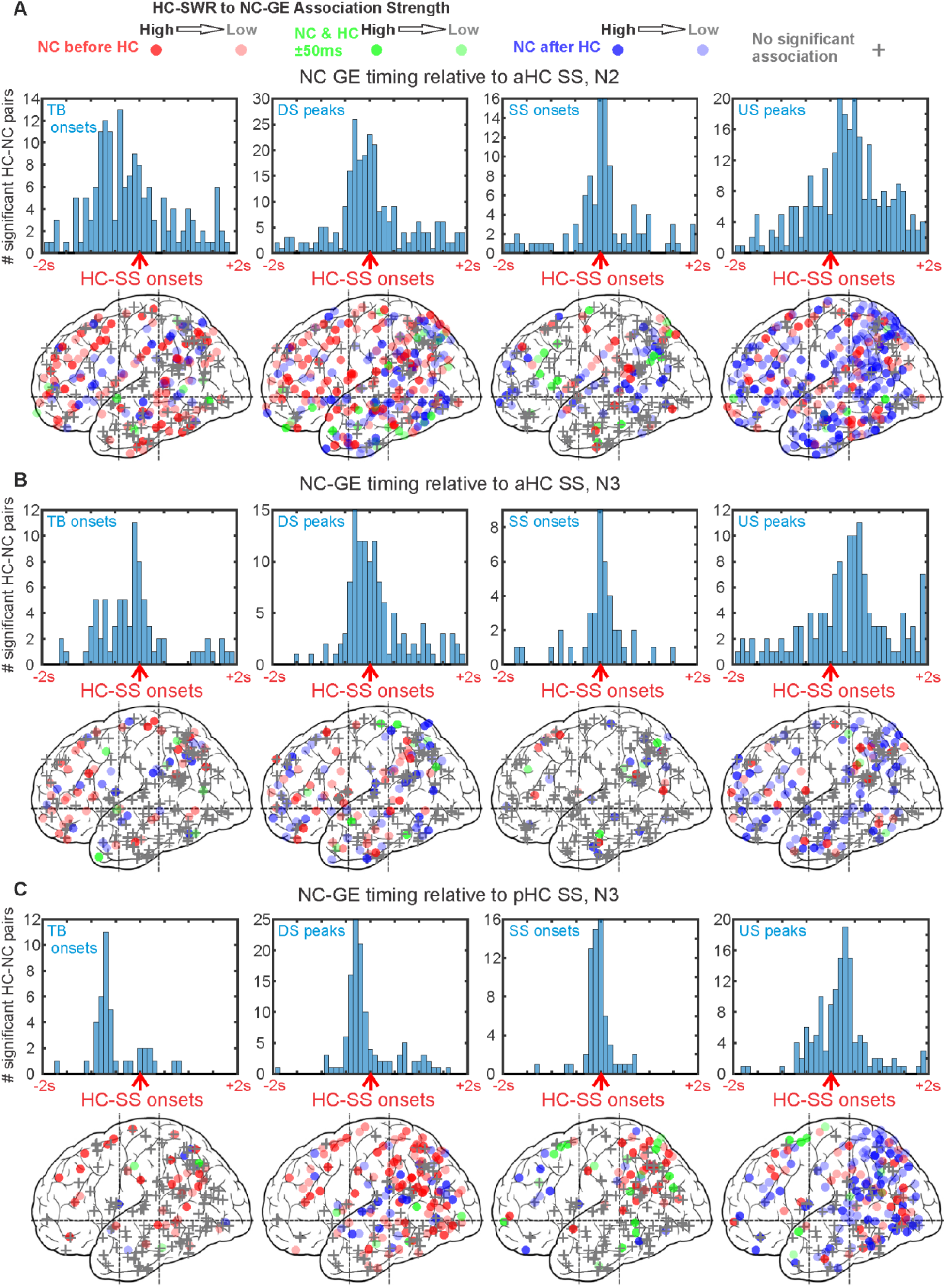
GEs across NC tend to occur with consistent latencies from HC-SS in N3. ***A***-***C***. Top rows: Histograms of peak latencies for HC-NC channel pairs with significant temporal correlations between HC-SS and NC-GE. Each count is the peak latency of a particular HC-NC channel pair. Bottom rows: maps of peak latency between HC-SS and NC-GE for cortical channels. Circles indicate where GE-SS relationships are significant, while plus signs represent non-significant channels. The intensity of each circle corresponds to the strength of NC-GE to HC-SS coupling estimated (as in Fig. 3), and the color of each circle indicates peak latency: red for NC-GE before HC-SS, blue for NC-GE after HC-SS, and green for NC-GE co-occurring with HC-SS (i.e. within 50 ms of each other). Both hemispheres and medial and lateral cortical sites are superimposed in each plot. ***A***. In N2, with aHC-SS as reference, NC-GE tend to occur in the following order: TB starts → DS peaks → SS starts → US peaks. ***B***. In N3, with aHC SS as reference, NC-GE tend to occur in the following order: DS peaks → SS starts → US peaks. ***C***. In N3, with pHC SS as reference, NC-GE tend to occur in the following order: TB starts → DS peaks → SS starts → US peaks. In ***A***-***C***, NC-SS tend to co-occur with HC-SS, and widespread NC-US tend to follow.

**Figure 4-3.**
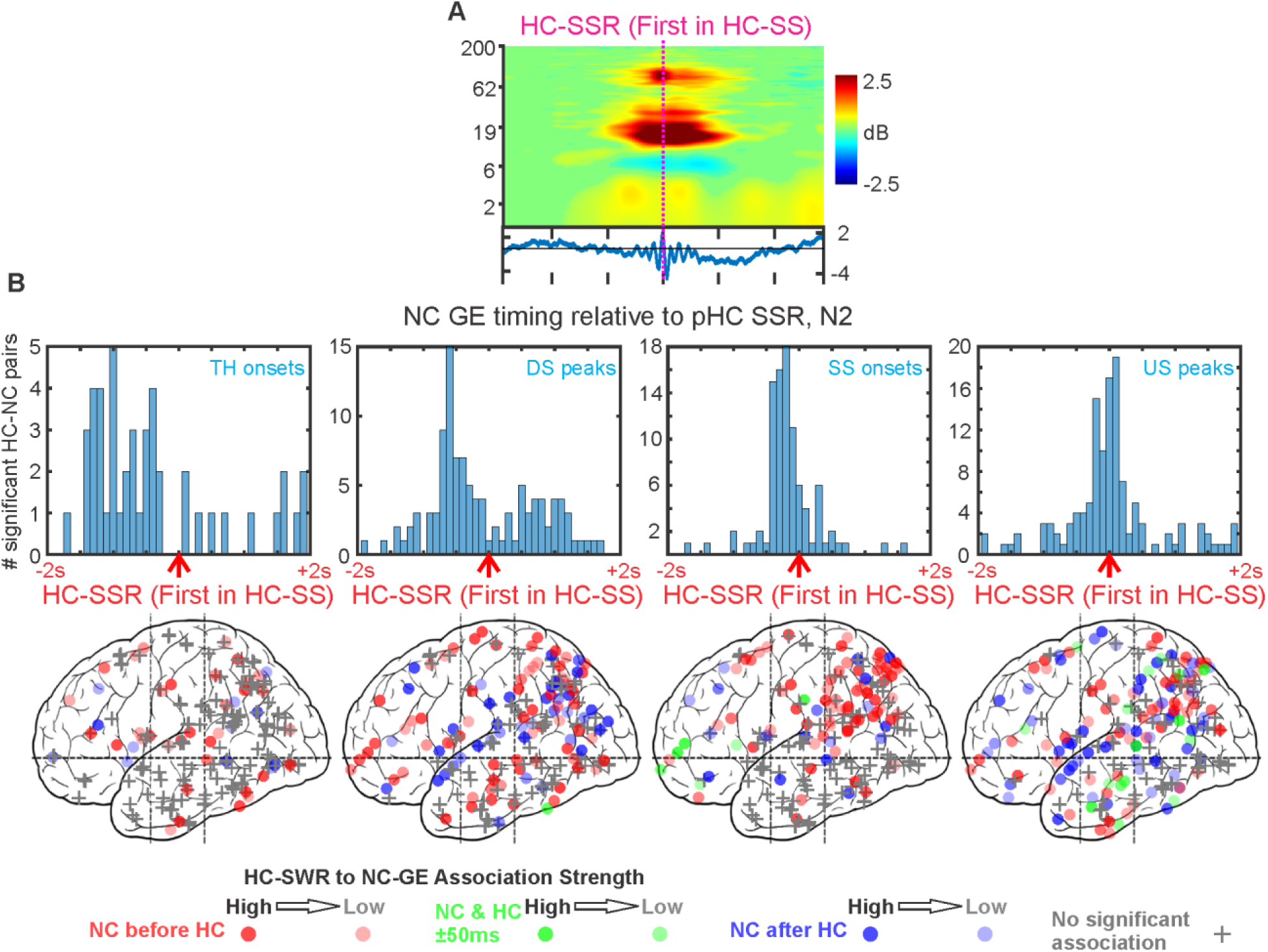
HC-SSR to NC-GE associations are preserved when limiting HC-SSR to only the first SSR within each HC-SS. ***A***. Event-related spectral power for first-in-SS HC-SSR (same patient source as Fig. 1D), with corresponding average LFP trace below. ***B***. Top row: Histograms of peak latencies for HC-NC channel pairs with significant temporal correlations between first-in-SS HC-SSR and NC-GE. Each count is the peak latency of a particular HC-NC channel pair. Bottom row: maps of peak latency between first-in-SS HC-SSR and NC-GE for cortical channels. Circles indicate where GE-SSR relationships are significant, while plus signs represent non-significant channels. The intensity of each circle corresponds to the strength of GE-SSR coupling estimated (as in Fig. 3), and the color of each circle indicates peak latency: red for NC-GE before HC-SSR, blue for NC-GE after HC-SSR, and green for NC-GE co-occurring with HC-SSR (i.e. within 50 ms of each other). Both hemispheres and medial and lateral cortical sites are superimposed in each plot. Similar to Fig. 4A, in N2, with pHC-SSR as reference, NC-GE tend to occur in the following order: TB starts → DS peaks → SS starts → US peaks.

**Figure 4-4.**
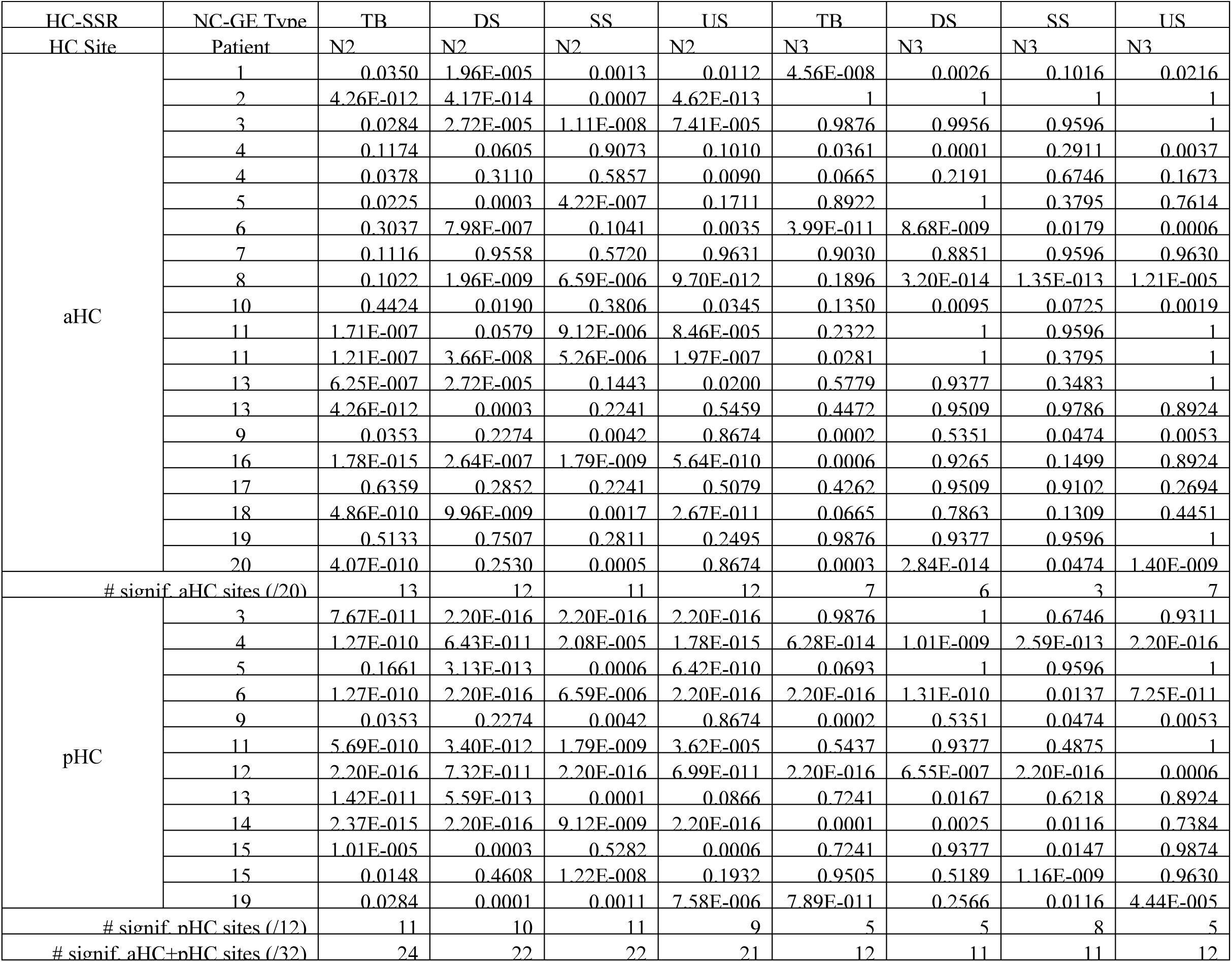
NC-GE synchronized with HC-SSR are not randomly distributed across sites. P-values for chi-square tests of homogeneity (α = 0.05 post FDR-correction applied over each column), conducted for each HC site with regard to each NC-GE type’s coupling to HC-SSR. Cells marked 2.20E-016 are placeholders for p < 2.2 × 10^-16^. # signif.: number of HC sites with a non-homogeneous anatomical distribution of significantly coupled NC sites. Out of 32 × 4 = 128 distributions of NC channels showing NC-GE to HC-SSR association in N2, 89 were significantly different from chance for HC-SSR in N2, compared to 46 in N3. The proportion of significant distributions was significantly greater in N2 (Fisher’s exact tests, p = 2.754 × 10^-8^). The percentages of pHC sites coupled to non-homogeneous NC site distributions (75-92% in N2, 42-67% in N3) were greater than the percentages of aHC sites (55-65% in N2, 15-35% in N3) for all NC-GE types.

**Figure 4-5.**
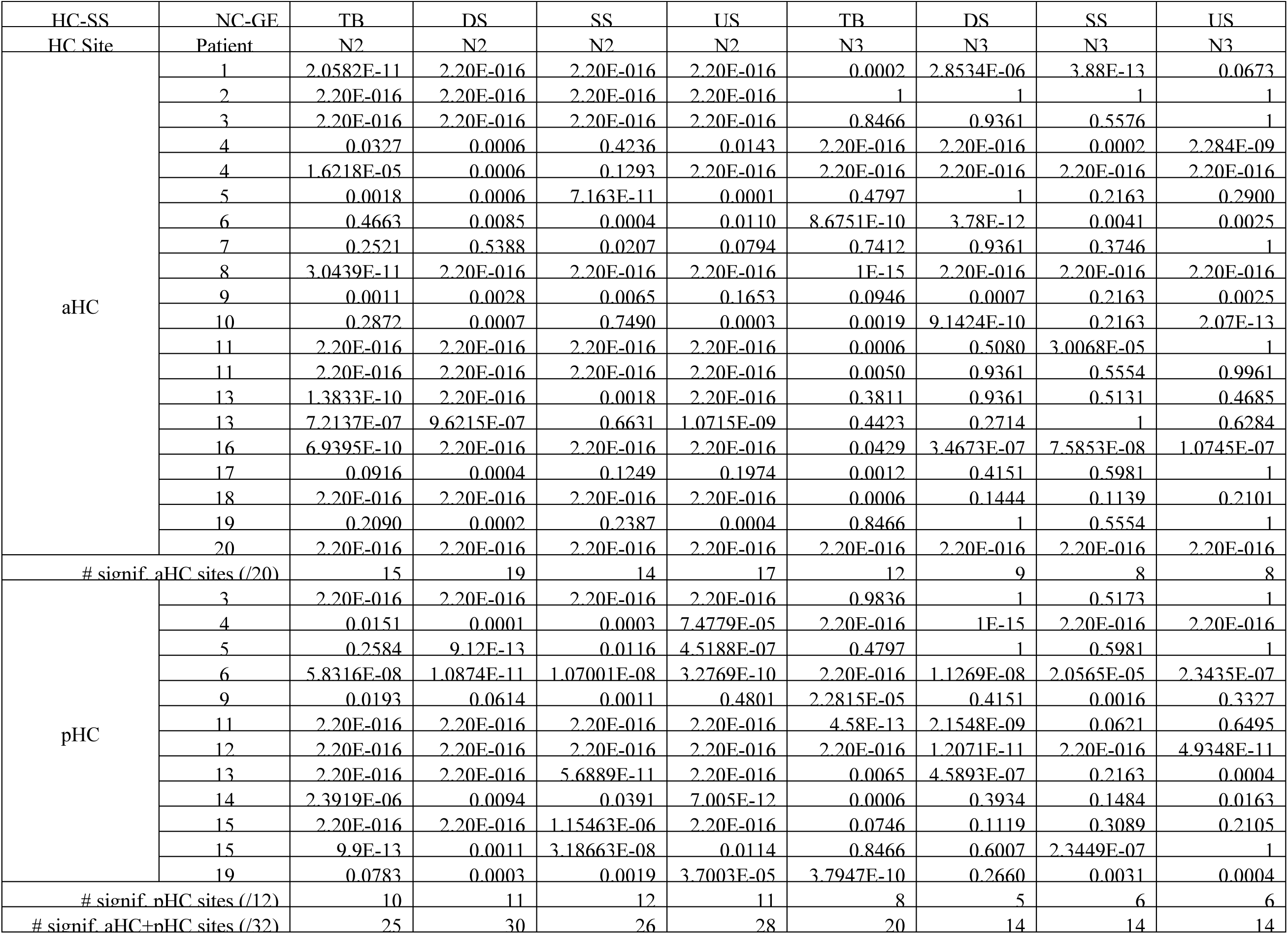
NC-GE synchronized with HC-SS are not randomly distributed across sites. P-values for chi-square tests of homogeneity (α = 0.05 post FDR-correction applied over each column), conducted for each HC site with regard to each NC-GE type’s coupling to HC-SS. Ant.: Ant-HC. Post: Post-HC. Cells marked 2.20E-016 are placeholders for p < 2.2 × 10^-16^. # signif.: number of HC sites with a non-homogeneous anatomical distribution of significantly coupled NC sites. Out of 32 × 4 = 128 distributions of NC channels showing NC-GE to HC-SS association in N2, 109 were significantly different from chance for HC-SSR in N2, compared to 62 in N3. The proportion of significant distributions was significantly greater in N2 (Fisher’s exact tests, p = 2.212 × 10^-10^). The pHC percentages (83-100% in N2, 42-67% in N3) and aHC percentages (70-95% in N2, 40-60% in N3) were both very high, consistent with a selective interaction with parietal areas.

**Figure 4-6.**
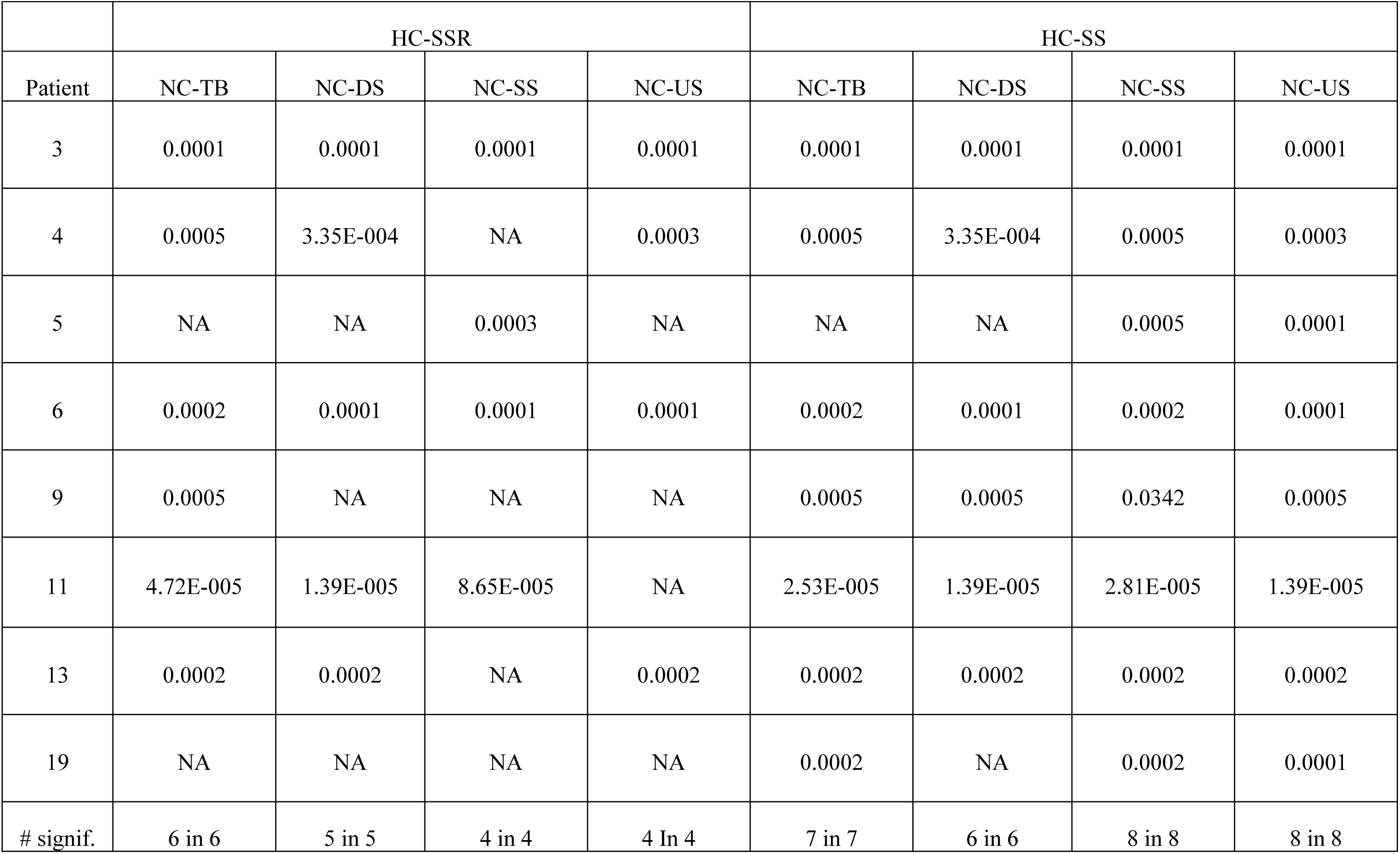
Coupled NC sites differ between simultaneously recorded aHC and pHC sites. Proportions of NC-GE from significantly coupled NC sites that overlapped with aHC-SSR/SS differ from the proportions for pHC-SSR/SS. Wilcoxon signed rank tests were performed on the data obtained for Figs. 4-4 and 4-5. Each test (whose p-value was tabulated above) required that both members of a HC site pair must have significant chi-square test result from Fig. 4-4 (for HC-SSR) or Fig. 4-5 (for HC-SS); otherwise, the corresponding cell would be filled with NA. # signif.: number of significant tests (in total number of qualified patients).

**Figure 6-1.**
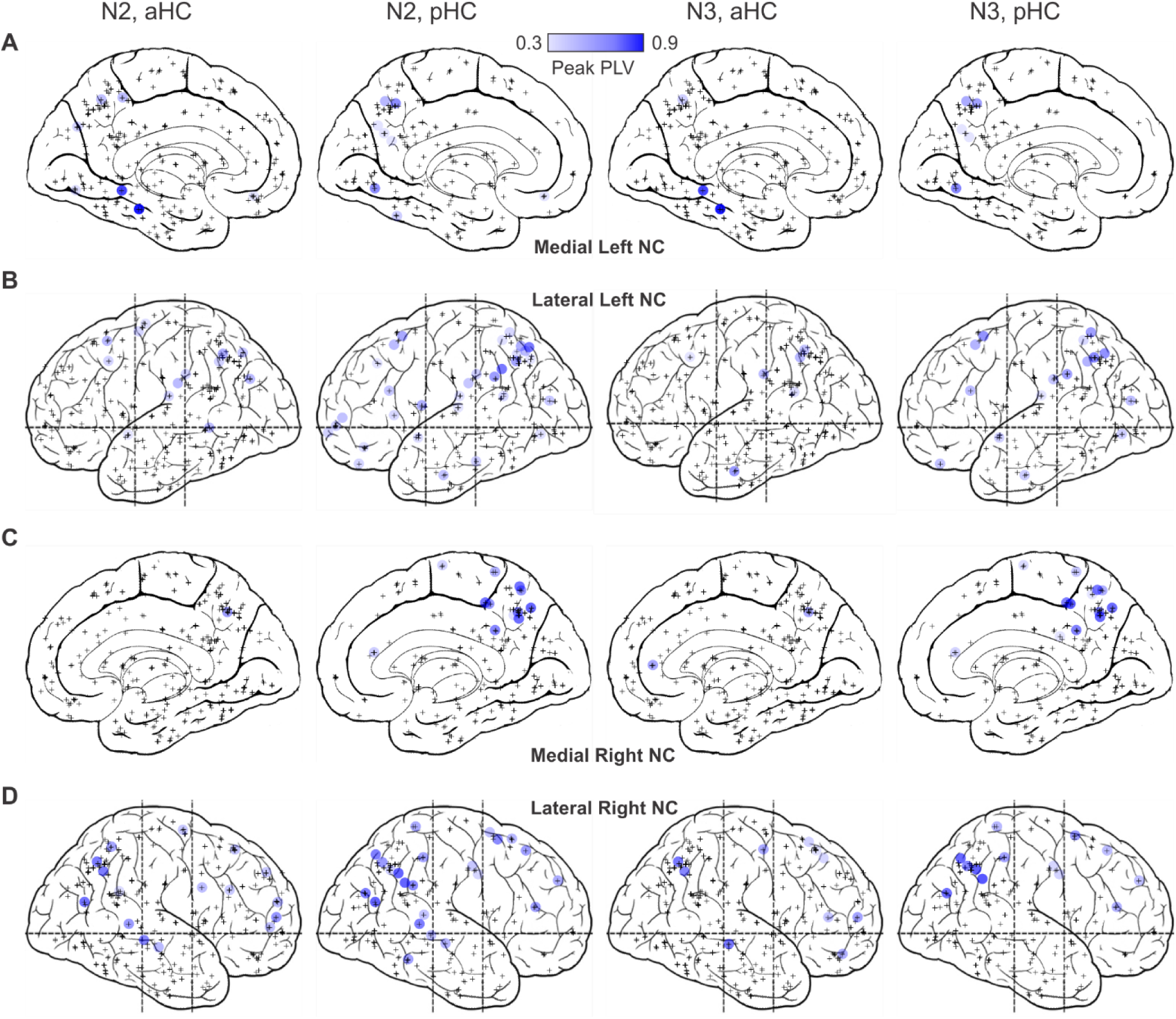
2d projection maps of left hemisphere medial (***A***), left hemisphere lateral (***B***), right hemisphere medial (***C***), and right hemisphere lateral (***D***) NC channels showing significant spindle phase-locking with HC. Each blue circle marks a significant NC channel, with color intensity corresponding to peak PLV amplitude. Plus signs mark channels with no significant phase-locking.

**Table 4-1.** ROIs for statistical analyses of spatio-temporal differences across NC regions in HC-NC relationships.

|  |  |
| --- | --- |
| Combined ROI | Original Freesurfer Label(s) |
| Lateral temporal | Banks of superior temporal gyrus, inferior temporal, middle temporal, superior temporal, temporal pole, transverse temporal |
| Cingulate | Caudal anterior cingulate, rostral anterior cingulate, isthmus cingulate, posterior cingulate |
| Orbitofrontal | Frontal pole, lateral orbitofrontal, medial orbitofrontal, pars orbitalis |
| Prefrontal | Caudal middle frontal, pars opercularis, pars triangularis, rostral middle frontal, superior frontal |
| Paracentral | Paracentral, postcentral, precentral |
| Medial temporo-occipital | Entorhinal, fusiform, lingual |
| Medial occipito-parietal | Cuneus, pericalcarine, precuneus |
| Insula | Insula |
| Lateral occipital | Lateral occipital |
| Lateral parietal | Inferior parietal, superior parietal, supramarginal |
| Fronto-central | All labels for the following Combined ROIs: Cingulate, orbito-frontal, prefrontal, paracentral |
| Non-frontal | All labels for the following Combined ROIs: Lateral temporal, medial temporo-occipital, medial occipito-parietal, insula, lateral occipital, lateral parietal |

